# JIP3 regulates bi-directional organelle transport in neurons through its interaction with dynein and kinesin-1

**DOI:** 10.1101/2021.10.11.463801

**Authors:** Ricardo Celestino, José B. Gama, Artur F. Castro-Rodrigues, Daniel J. Barbosa, Ennio A. d’Amico, Andrea Musacchio, Ana Xavier Carvalho, João H. Morais-Cabral, Reto Gassmann

## Abstract

The conserved MAP kinase and motor scaffold JIP3 prevents excess lysosome accumulation in axons of vertebrates and invertebrates. Whether and how JIP3’s interaction with dynein and kinesin-1 contributes to this critical organelle clearance function is unclear. Using purified recombinant human proteins, we show that dynein light intermediate chain (DLIC) binds to the N-terminal RH1 domain of JIP3, its paralog JIP4, and the lysosomal adaptor RILP. A point mutation in a hydrophobic pocket of the RH1 domain, previously shown to abrogate RILPL2 binding to myosin Va, abrogates the binding of JIP3/4 and RILP to DLIC without perturbing the interaction between the JIP3 RH1 domain and kinesin heavy chain. Characterization of this separation-of-function mutation in *Caenorhabditis elegans* shows that JIP3–bound dynein is required for organelle clearance in the anterior process of touch receptor neurons. Unlike JIP3 null mutants, JIP3 that cannot bind DLIC causes prominent accumulation of endo-lysosomal organelles at the neurite tip, which is rescued by a disease-associated point mutation in JIP3’s leucine zipper that abrogates kinesin light chain binding. These results highlight that RH1 domains are interaction hubs for cytoskeletal motors and suggest that JIP3–bound dynein and kinesin-1 participate in bi-directional organelle transport.

## INTRODUCTION

Long-range intracellular transport of vesicles and organelles by microtubule-based motors is critical for the development, survival, and function of neurons (Guedes-Dias and Holzbaur, 2019). In the axon, where most microtubules assume a plus-end-out orientation, multiple kinesins mediate anterograde transport toward synapses, and dynein is responsible for retrograde transport toward the cell body. How kinesins and dynein work together in axons to form a useful bi-directional transport system is a fundamental unanswered question.

c-Jun N-terminal kinase-interacting protein 3 (JIP3), also known as Sunday Driver or JSAP1, has emerged as an evolutionarily conserved regulator of axonal transport that associates with both kinesin-1 and dynein. JIP3 was originally identified as a scaffolding protein that binds c-Jun N-terminal kinase (JNK) and kinesin light chain (KLC) (Bowman *et al*., 2000; Ito *et al*., 1999; Kelkar *et al*., 2000; Verhey *et al*., 2001), and as the product of the *sunday driver* (*syd*) gene in a *D. melanogaster* screen for axonal transport mutants (Bowman *et al*., 2000). The *syd* mutants accumulate synaptic vesicle cargo in segmental nerve axons, a defect that is also observed in *kinesin heavy chain (khc)* and *klc* mutants (Bowman *et al*., 2000). Similarly, mutants of *C. elegans unc:16/JIP3, unc:116/KHC, and jnk:1/JNK* mis-localize synaptic vesicles to the dorsal processes of cholinergic motor neurons (Byrd *et al*., 2001), and *unc:16* mutants are defective in the sorting of synaptic vesicle cargo at the trans golgi (Choudhary *et al*., 2017). In addition to regulating composition and distribution of golgi-derived transport vesicles, work in *C. elegans* established that UNC-16 acts as a negative regulator of axonal organelle abundance: mitochondria, lysosomes, early/recycling endosomes, and the golgi all accumulate in cholinergic motor neuron axons of *unc:16* mutants (Edwards *et al*., 2013; Edwards *et al*., 2015), suggesting that one key function of UNC-16 is to clear organelles from axons. Studies in zebrafish sensory neurons, in mouse embryo cortical neurons, and in human iPSC-derived neurons have since shown that JIP3 also negatively regulates axonal lysosome abundance in vertebrates (Drerup and Nechiporuk, 2013; Gowrishankar *et al*., 2017; Gowrishankar *et al*., 2021), and that JIP3 shares this function with its paralog JIP4 (Gowrishankar *et al*., 2021). Recent work identified missense mutations in the human JIP3- encoding gene MAPK8IP that cause neurodevelopmental disorders and intellectual disability (Iwasawa *et al*., 2019; Platzer *et al*., 2019), and modelling in *C. elegans* showed that some of these mutations impair JIP3’s organelle clearance function (Platzer *et al*., 2019).

How JIP3’s association with microtubule-based opposite-polarity motors contributes to its role as a negative regulator of axonal organelle abundance remains poorly explored. Most studies addressing JIP3’s motor-dependent functions have focused on its interaction with kinesin-1, which is a heterotetramer composed of a KHC homodimer and a KLC homodimer. The structure of the KLC tetratricopeptide repeat (TPR) domain bound to the mouse JIP3 leucine zipper has been determined by X-ray crystallography (Cockburn *et al*., 2018), and one of the recently described disease mutations in human JIP3 is predicted to interfere with KLC binding (Platzer *et al*., 2019). JIP3 also binds the C-terminal tail of KHC, and pull-downs of JIP3 fragments from mouse brain lysate mapped the KHC binding site to the JIP3 N-terminus (Sun *et al*., 2011). Studies in cultured neurons with JIP3 mutants harboring deletions in the KHC and/or KLC binding region suggest that the interaction with kinesin-1 stimulates axonal elongation (Muresan and Muresan, 2005; Sun *et al*., 2011; Sun *et al*., 2013; Watt *et al*., 2015). Furthermore, both the KHC and the KLC binding region of JIP3 have been implicated in the activation of kinesin-1 motility (Sato *et al*., 2015; Sun *et al*., 2011; Sun *et al*., 2017; Watt *et al*., 2015). Besides JIP3 itself (Byrd *et al*., 2001; Verhey *et al*., 2001), the only firmly established cargo of JIP3–bound kinesin-1 is the BDNF receptor TRKB, which is transported from the cell body to synapses in the distal axon (Drerup and Nechiporuk, 2013; Huang *et al*., 2011; Ma *et al*., 2017; Sun *et al*., 2017). Additionally, a study in mouse hippocampal neurons reported slowed anterograde transport of mitochondria and amyloid precursor protein in a JIP3 mutant defective for kinesin-1 binding (Sato *et al*., 2015). There is also evidence that JIP3 and the structurally unrelated JIP1 (another MAP kinase scaffold that associates with kinesin-1 and dynein) cooperate during anterograde transport: in differentiated CAD cells, low-level overexpression of either adaptor stimulates localization of the other adaptor to the neurite tip (Hammond *et al*., 2008), and overexpression of either adaptor in mouse hippocampal neurons enhances TRKB localization to the axonal tip (Sun *et al*., 2017).

Although JIP3 has been primarily characterized as a kinesin-1 adaptor, the fact that JIP3 inhibition causes accumulation of vesicles and organelles in axons implies a role in promoting retrograde transport. The first direct evidence for such a role came from a zebrafish study, which demonstrated that retrograde motility of activated JNK and lysosomes is impaired in JIP3 mutant axons (Drerup and Nechiporuk, 2013). More recently, JIP3 inhibition was shown to impair retrograde autophagosome motility in the *C. elegans* AYI interneuron and in rat hippocampal neurons (Cason *et al*., 2021; Hill *et al*., 2019). Consistent with a role for JIP3 in retrograde transport, *C. elegans* dynein light intermediate chain (DLIC) can pull down UNC- 16 from COS-7 cell lysate, albeit only when the two proteins are co-expressed together with KLC (Arimoto *et al*., 2011); and JIP3’s leucine zipper region can pull down the dynein co-factor dynactin from HeLa cell lysate (Montagnac *et al*., 2009). The molecular details and functional significance of these interactions remain to be determined. Moreover, it is unclear whether JIP3–bound kinesin-1 and JIP3–bound dynein act on the same type of cargo, or whether JIP3 associates with the two motors in separate to carry out distinct functions, namely anterograde transport of vesicle cargo through kinesin-1 and axonal organelle clearance through dynein (Miller, 2017).

JIP3 and its vertebrate paralog JIP4 share their overall domain organization with proteins of the Rab-interacting lysosomal protein (RILP) family, which in vertebrates includes RILP, RILP-like 1 (RILPL1), and RILPL2 (Vilela *et al*., 2019; Wang *et al*., 2004). RILP recruits dynein-dynactin to late endosomes and lysosomes (Jordens *et al*., 2001), and RILPL2 is a cargo adaptor for myosin Va involved in ciliogenesis (Lisé *et al*., 2009; Schaub and Stearns, 2013). RILPL1 also plays a role in ciliogenesis (Schaub and Stearns, 2013), but an interaction between RILPL1 and cytoskeletal motors has yet to be demonstrated. The homodimeric RILP/JIP3 superfamily is characterized by an N-terminal RILP homology (RH) 1 domain of approximately 100 residues, which consists of a four-helix bundle followed by a short coiled-coil segment (Wei *et al*., 2013); by a coiled-coil region that extends C-terminally from the RH1 domain; and by a four-helix RH2 domain located at variable distance from the RH1 domain (Fig. 1 A). RILPL2’s RH1 domain binds to myosin Va (Wei *et al*., 2013), JIP3’s RH1 domain includes the KHC-interacting region (Sun *et al*., 2011), and the RH2 domain binds small GTPases of the Rab family (Wang *et al*., 2004; Wu *et al*., 2005). Interestingly, residues 1-240 of *C. elegans* UNC-16 interact with DLI-1/DLIC in the yeast 2-hybrid assay (Arimoto *et al*., 2011), raising the question of whether the KHC binding site at the JIP3 N-terminus overlaps with that of DLIC.

**Figure 1.**
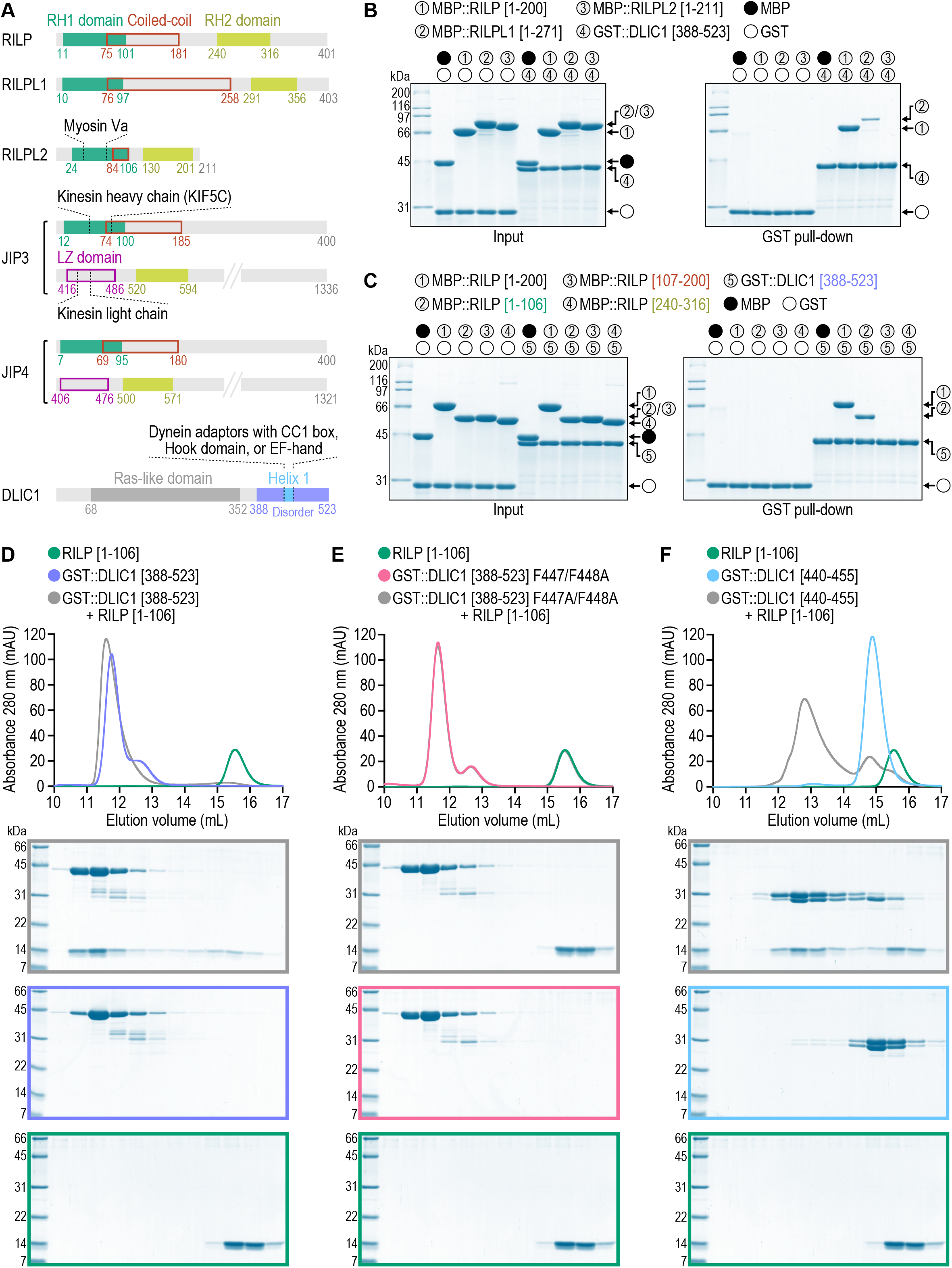
Dynein adaptors of the RILP/JIP3 superfamily use their N-terminal RH1 domain to bind the C-terminal helix 1 of dynein light intermediate chain. **(A)** Domain organization of human dynein light intermediate chain 1 (DLIC1) and the 5 human cargo adaptors for cytoskeletal motors characterized by the presence of RILP homology (RH) 1 and RH2 domains. JIP3 and JIP4 also have a second coiled-coil region, here called the leucine zipper (LZ) domain. Residue numbers are from UniProt entries Q96NA2-1 (RILP), Q5EBL4-1 (RILPL1), Q969X0-1 (RILPL2), Q9UPT6-1 (JIP3), O60271-1 (JIP4), and Q9Y6G9-1 (DLIC1). (B), (C) *(left)* Coomassie Blue-stained SDS- PAGE gel of purified recombinant protein mixtures prior to addition of glutathione agarose resin (Input). *(right)* Coomassie Blue-stained SDS-PAGE gel of proteins eluted from glutathione agarose resin after GST pull-down. Proteins correspond to the human homologs described in *(A)* except for RILPL1, which is the mouse homolog (Q9JJC6-1). Molecular weight is indicated in kilodaltons (kDa). **(D) - (F)** Elution profiles *(top)* and Coomassie Blue-stained SDS-PAGE gels *(bottom)* of purified recombinant human proteins after size exclusion chromatography on a Superdex 200 Increase column. Molecular weight is indicated in kilodaltons (kDa).

Here we show that the RH1 domains of RILP, RILPL1, and JIP3/4 are receptors for a conserved C-terminal helix in DLIC that is known to bind several functionally and structurally distinct dynein adaptors (Celestino *et al*., 2019; Lee *et al*., 2018; Lee *et al*., 2020; Renna *et al*., 2020). We show that the interaction occurs through a hydrophobic pocket in the RH1 domain that in RILPL2 binds Myosin Va, and we validate a point mutation in the RH1 domain that specifically abrogates binding to DLIC without interfering with KHC binding by JIP3. Functional characterization of this JIP3 separation-of-function mutant in *C. elegans* demonstrates that UNC-16/JIP3’s interaction with DLIC promotes retrograde transport of endo-lysosomal organelles and synaptic vesicles in the anterior process of touch receptor neurons, which have an axon-like microtubule organization. Characterization of additional engineered mutants, including a disease-associated UNC-16 mutation that abrogates KLC binding, supports the idea that JIP3 uses kinesin-1 and dynein to drive bi-directional transport of endo-lysosomal organelles.

## RESULTS

### Dynein light intermediate chain’s C-terminal helix 1 binds to the RH1 domains of RILP, JIP3, and JIP4

Vertebrates express 5 proteins with RH1/2 domains: JIP3 and its paralog JIP4, and the three members of the RILP family (Fig. 1 A). RILP and JIP3 are known to bind dynein light intermediate chain (DLIC), yet they contain none of the three previously characterized structural elements that other cargo adaptors use to accommodate the amphipathic helix 1 in the DLIC C-terminal tail (DLIC-C). Whether JIP4 and the other two members of the RILP family, RILPL1 and RILPL2, interact with DLIC-C has not been tested. We therefore set out to examine how DLIC-C interacts with RH1/2-containing proteins using purified recombinant human proteins (except for mouse RILPL1). Pull-downs of GST-tagged DLIC1[388-523], which corresponds to DLIC-C of isoform 1, showed that DLIC1-C interacts with the N-terminal halves of RILP and RILPL1, while no interaction was detectable with full-length RILPL2 (Fig. 1 B; note that RILPL2 is half the size of RILP/RILPL1). Pull-downs with additional RILP fragments showed that RILP[1-106], which corresponds to the RH1 domain (RILP-RH1), is sufficient to bind DLIC1-C (Fig. 1 C). Neither the subsequent coiled-coil region RILP[107-200] nor the RH2 domain RILP[240-316] bound to DLIC1-C (Fig. 1 C). We confirmed the interaction between RILP-RH1 and DLIC1-C by size exclusion chromatography (SEC) (Fig. 1 D). Further SEC experiments showed that introducing two point mutations (F447A/F448A) into helix 1 of DLIC1-C abrogates binding to RILP-RH1, and that DLIC1[440-455], which corresponds to helix 1, is sufficient for binding (Fig. 1, E and F). Analogous experiments with the RH1 domains of JIP3 and JIP4 confirmed that they also bind DLIC1-C, and that they do so in a manner that depends on DLIC1-C helix 1 (Fig. 2; Fig. 3; and Fig. S2). The RH1 domain therefore joins the Hook domain, the CC1 box, and the EF-hand as a fold that can accommodate helix 1 of the dynein light intermediate chain C-terminal tail (Lee *et al*., 2018; Celestino *et al*., 2019; Lee *et al*., 2020).

**Figure 2.**
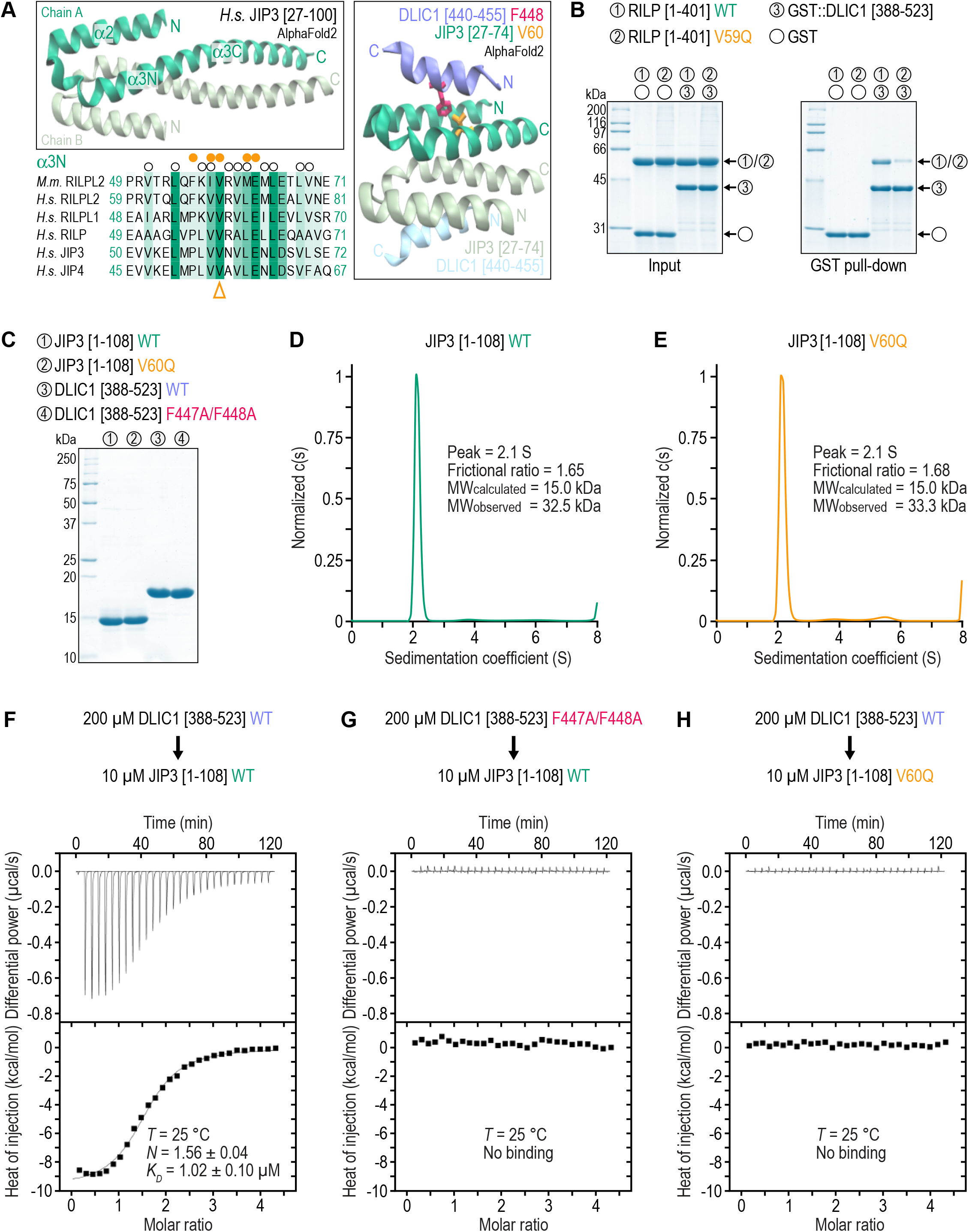
The C-terminal amphipathic helix 1 of dynein light intermediate chain binds the RH1 domain’s hydrophobic pocket. **(A)** *(top left)* Model of the dimeric RH1 domain of human JIP3 (UniProt entry Q9UPT6- 1) generated using the ColabFold Jupyter Notebook for protein structure prediction (Mirdita *et al*., 2021). *(bottom left)* Sequence alignment of the α3N helix for the RILP/JIP3 superfamily. Open circles denote residues participating in dimer formation in mouse RILPL2 (Wei *et al*., 2013). Closed circles denote residues participating in the interaction between RILPL2 and myosin Va. Arrowhead points to the valine residue that forms part of the RH1 domain’s hydrophobic pocket and whose mutation to glutamine in RILPL2 abrogates the interaction with myosin Va. *(right)* Model of the JIP3 RH1 domain’s 4-helix bundle with the C-terminal helix 1 of DLIC1 (residues 440-455) docked at the hydrophobic pocket, as predicted by ColabFold. A phenylalanine and a valine side chain in DLIC1 and JIP3, respectively, which we mutate in this study, are also rendered. The models have a per-residue confidence (pLDDT) of > 90 % throughout and a consistently low alignment error (PAE) (Tunyasuvunakool *et al*., 2021). **(B)** *(left)* Coomassie Blue-stained SDS-PAGE gel of purified recombinant protein mixtures prior to addition of glutathione agarose resin (Input). *(right)* Coomassie Blue-stained SDS-PAGE gel of proteins eluted from glutathione agarose resin after GST pull-down. Proteins correspond to the human homologs. WT denotes wild-type. Molecular weight is indicated in kilodaltons (kDa). **(C)** Coomassie Blue-stained SDS-PAGE gel of purified recombinant human proteins used in AUC and ITC experiments. Molecular weight is indicated in kilodaltons (kDa). **(D), (E)** Sedimentation velocity AUC profiles with theoretical (MW_calculated_) and experimentally measured molecular mass (MW_observed_). The MW_observed_ values indicate that both proteins are dimeric in solution. **(F) - (H)** Thermograms and binding isotherms of representative ITC titrations. JIP3[1-108] concentration is the concentration of the dimer. The dissociation constant (*K_D_*) and the binding stoichiometry (*N*) are given as mean ± SD (n=3) and were derived from fitting to the binding isotherm (black line) with a One Set of Sites model using ORIGIN software.

**Figure 3.**
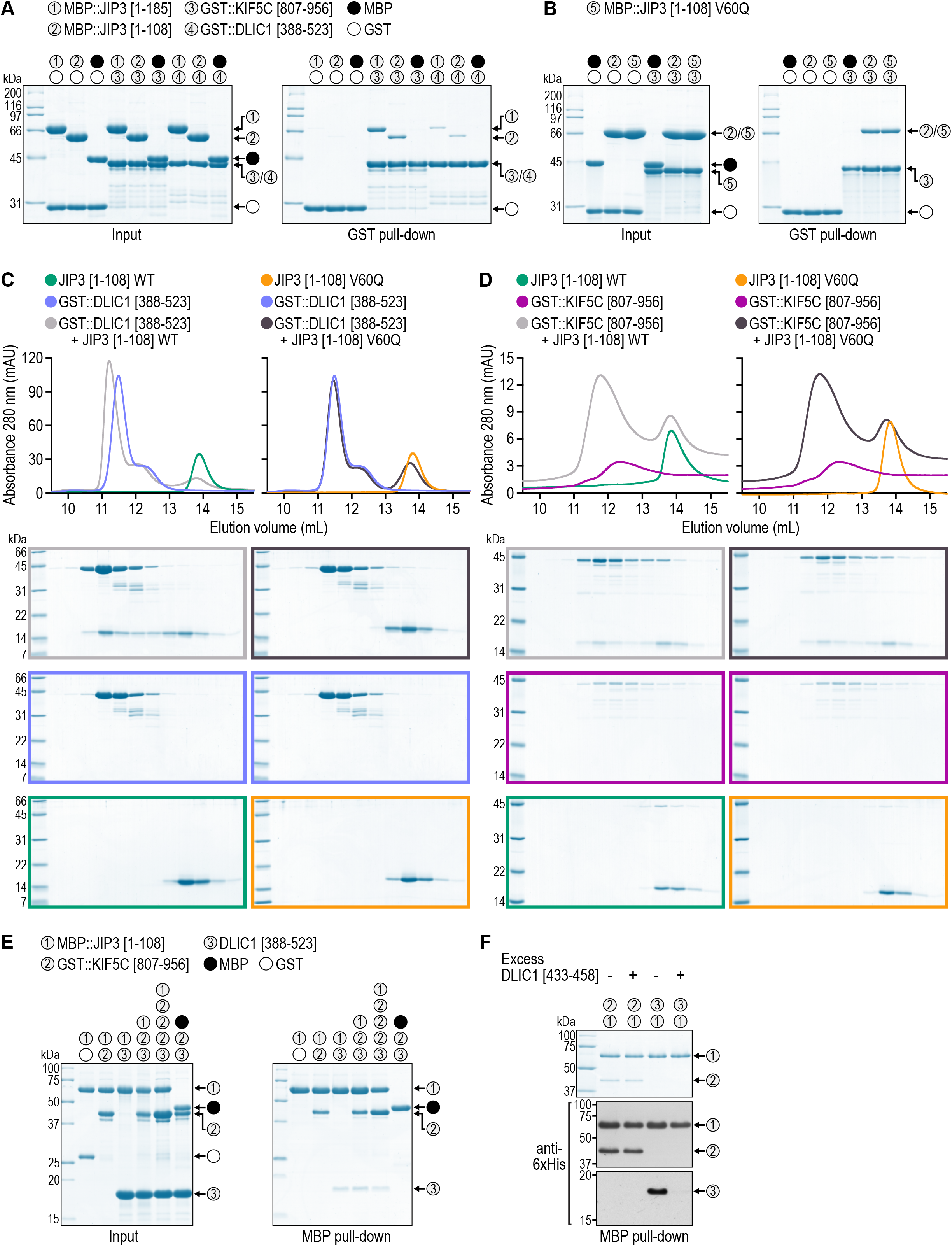
Kinesin heavy chain also binds to the RH1 domain of JIP3 but does not compete for binding with dynein light intermediate chain. **(A), (B)** *(left)* Coomassie Blue-stained SDS-PAGE gel of purified recombinant protein mixtures prior to addition of glutathione agarose resin (Input). *(right)* Coomassie Blue-stained SDS-PAGE gel of proteins eluted from glutathione agarose resin after GST pull-down. Proteins correspond to the human homologs described in Fig. 1A and to mouse kinesin heavy chain KIF5C (UniProt entry P28738-1). Molecular weight is indicated in kilodaltons (kDa). **(C), (D)** Elution profiles *(top)* and Coomassie Blue-stained SDS-PAGE gels *(bottom)* of purified recombinant human proteins after size exclusion chromatography on a Superdex 200 Increase column. WT denotes wild-type. Molecular weight is in kilodaltons (kDa). (E) *(left)* Coomassie Blue-stained SDS-PAGE gel of purified recombinant protein mixtures prior to addition of amylose resin (Input). *(right)* Coomassie Blue-stained SDS-PAGE gel of proteins eluted from amylose resin after MBP pull-down. The actual amount of DLIC1[388-523] in the pull down reaction was 5-fold higher than what is shown for the input. The KIF5C fragment was used at two concentrations that differ 3-fold, as indicated above the 5th and 6th lanes from the left. **(F)** Coomassie Blue-stained SDS-PAGE gel *(top)* and corresponding immunoblot *(bottom)* of proteins eluted from amylose resin after MBP pull-down as in *(E)*. All proteins contain a 6xHis tag (see Materials and Methods) that is detected on the immunoblot. Proteins are the same as in *(E)*, but amounts in the pull-down mixture were decreased relative to those in *(E)* such that DLIC1-C helix 1 peptide could be added in 150-fold molar excess over the KIF5C fragment.

### Dynein light intermediate chain binding occurs via a hydrophobic pocket in the RH1 domain

Our pull-down experiments suggested that all RH1/2 domain-containing proteins except RILPL2 are adaptors for dynein. This is consistent with the well-established role of RILPL2 as an adaptor for myosin Va (Lisé *et al*., 2009; Wei *et al*., 2013). Structural analysis showed that the globular tail domain of myosin Va interacts with the RH1 domain of RILPL2 in part by inserting a short helical segment into a hydrophobic pocket that is formed by the RH1 helical bundle, with two binding sites per RH1 dimer (Wei *et al*., 2013) (Fig. 2 A). Since the amphipathic DLIC-C helix 1 is known to insert into hydrophobic pockets present in the Hook domain, CC1 box, and EF-hand of different adaptors (Lee *et al*., 2018; Lee *et al*., 2020), we speculated that the hydrophobic pocket of the RH1 domain might be used to accommodate DLIC-C helix 1 in RILP, RILPL1, JIP3, and JIP4. High-confidence structure predictions, generated by the AlphaFold2-based pipeline ColabFold (Mirdita *et al*., 2021), suggest that this is indeed the case (Fig. 2 A). Mutating a single residue (V59Q) in the hydrophobic pocket of RILPL2-RH1 abrogates the myosin Va interaction (Wei *et al*., 2013). To directly test whether the other RH1 domains bind DLIC-C helix 1 through the same hydrophobic pocket, we introduced the analogous point mutation into human RILP (V59Q), JIP3 (V60Q), and JIP4 (V55Q) (Fig. 2 A). GST pull-downs with purified recombinant proteins showed that full-length RILP carrying the V59Q mutation had significantly reduced affinity for GST-tagged DLIC1-C (Fig. 2 B), as did mutated RILP-RH1 in SEC experiments (Fig. S1). To gain a more quantitative understanding of the interaction, we sought to perform isothermal titration calorimetry (ITC). For RILP[1-106], analytical ultracentrifugation (AUC) experiments indicated that the protein was present as a mixture of dimer and trimer (data not shown). Because of this heterogeneity of RILP-RH1, we used JIP3[1-108] and JIP4[1-103] for ITC experiments, since AUC runs showed that these proteins behave as monodisperse dimers (Fig. 2, C and D; and Fig. S2, A and B). ITC measurements revealed that JIP3-RH1 and JIP4-RH1 bind to DLIC1-C with low micromolar affinity (K_D_ of ∼1 µM and ∼4 µM, respectively) and with a reaction stoichiometry that indicates two DLIC1-C molecules per RH1 dimer (Fig. 2 F and Fig. S2 C). This is similar to what has been determined for the interaction between DLIC1-C and other dynein adaptors (Lee *et al*., 2018; Lee *et al*., 2020). As expected, there was no detectable signal in the thermogram when DLIC1-C helix 1 was mutated (Fig. 2 G and Fig. S2 D). Strikingly, introducing the V/Q mutation into JIP3-RH1 and JIP4-RH1 had the same effect (Fig. 2 H and Fig. S2 E). Importantly, AUC confirmed that the V60Q mutation does not perturb dimerization of JIP3-RH1 (Fig. 2 E), consistent with previous analysis of this mutation in RILPL2 (Wei *et al*., 2013). These results suggest that RILP, JIP3, JIP4, and most likely also RILPL1, bind to DLIC1-C helix 1 via a hydrophobic pocket in their RH1 domain, which in RILPL2 accommodates helix 2 of myosin Va’s globular tail domain.

### The JIP3 V60Q mutation does not affect binding to kinesin heavy chain

JIP3 binds the C-terminal tail of KHC, and pull-downs from mouse brain lysate using JIP3 fragments previously suggested that residues 50-80 of JIP3 are required for the interaction (Sun *et al*., 2011). Since the JIP3 V60Q mutation falls within this region, we asked whether this mutation also affects binding to the KHC tail. Using purified recombinant proteins, we first confirmed that JIP3-RH1 binds GST-tagged mouse KIF5C[807-956] using GST pull-downs and SEC (Fig. 3, A and D). Although we could not perform ITC measurements with KIF5C[807-956] because of the poor solubility of this fragment, a side-by-side comparison of pull-downs indicated that the KHC tail binds more strongly to JIP3-RH1 than DLIC1-C (Fig. 3 A). Introducing the V60Q mutation into JIP3-RH1 abrogated binding to DLIC1-C but had no effect on the interaction with the KHC tail in GST pull-down and SEC experiments (Fig. 3, B - D). Of note, the amount of GST::KIF5C[807-956] eluting from the SEC column increased significantly when JIP3-RH1 was present, suggesting that binding to the RH1 domain improves the solubility of the KHC tail fragment (Fig. 3 D). The JIP3-RH1 V60Q mutant had the same stabilizing effect on GST::KIF5C[807-956], which further supports the idea that this mutation does not perturb the interaction with the KHC tail. We conclude that JIP3-RH1 binds to both DLIC and KHC, and that JIP3 V60Q is a separation-of-function mutation that specifically abrogates the interaction between JIP3 and DLIC.

We also sought to address whether the KHC tail and DLIC1-C can bind to JIP3-RH1 simultaneously. Unfortunately, at the relatively low protein concentrations we had to use for SEC experiments with the KHC tail (which exhibited poor solubility even when tagged with GST), JIP3-RH1 and DLIC1-C no longer co-eluted robustly from the column, so we could not use SEC to determine whether JIP3-RH1 can form a tripartite complex with the KHC tail and DLIC1-C. As an alternative approach, we performed pull downs with MBP-tagged JIP3[1-108] in the presence of GST::KIF5C[807-956] alone, DLIC1[388-523] alone, or both GST::KIF5C[807-956] and DLIC1[388-523]. The KHC tail and DLIC1-C were pulled down by JIP3-RH1 in similar amounts regardless of whether or not the other fragment was present (Fig. 3 E). To further probe whether the KHC tail and DLIC1-C compete for binding to JIP3-RH1, we added an excess of DLIC-C helix 1 peptide (residues 433-458) to the pull down mixture. This displaced DLIC-1-C but not the KHC tail from JIP3-RH1 (Fig. 3 F). These experiments suggest that significant overlap between the DLIC and KHC binding sites on JIP3-RH1 is unlikely.

### *C. elegans* UNC-16 V72Q is equivalent to human JIP3 V60Q

To explore the functional significance of the interaction between DLIC and JIP3, we turned to *C. elegans*, whose *unc:16* gene encodes the single homolog of JIP3 and JIP4. *C. elegans* offers the advantage that *unc:16* mutants can be propagated in a homozygous state, so there is no residual maternal wild-type protein which could potentially confound phenotypic interpretation. Before proceeding to *in vivo* analysis, we asked whether the interactions we identified among human proteins are conserved in the nematode. Using pull-down assays with purified recombinant proteins, we confirmed that UNC-16[1-120], which corresponds to the RH1 domain, binds to DLI-1[369-442], which corresponds to DLIC-C; and that the V72Q mutation in UNC-16, which corresponds to the V60Q mutation in human JIP3, abrogates binding to DLIC-C (Fig. S3 A). Furthermore, UNC-16[1-120] bound to UNC-116[675-815], which corresponds to the KHC tail, and the V72Q mutation in UNC-16 had no apparent effect on this interaction (Fig. S3 B). We conclude that human JIP3 V60Q and *C. elegans* UNC-16 V72Q are equivalent mutations: they abrogate RH1 domain binding to DLIC without perturbing the interaction between the RH1 domain and KHC.

### Dynein light intermediate chain binding is essential for UNC-16 function *in vivo*

We next generated animals expressing UNC-16 V72Q by editing the *unc:16* locus using the CRISPR/Cas9 method (Fig. 4 A). To assess UNC-16 protein levels, we generated an affinity-purified antibody, which recognizes a protein of the predicted size for full-length UNC-16 (134 kDa) on immunoblots of adult worm lysate (Fig. 4 B). This protein is missing in animals carrying *unc:16(ce483)*, a previously characterized loss-of-function mutation (Edwards *et al*., 2013). Immunoblotting confirmed that UNC-16 V72Q is expressed at the same level as wild-type UNC-16 (Fig. 4 B). To assess the effect of the *unc:16(V72Q)* mutation on animal behavior, we determined body bending frequency in liquid at the young adult stage. This showed that *unc:16(V72Q)* animals are locomotion-deficient, and that this phenotype is as severe as in *unc:16(ce483)* animals (Fig. 4 C). We conclude that the interaction with DLI- 1 is essential for UNC-16 function *in vivo*.

**Figure 4.**
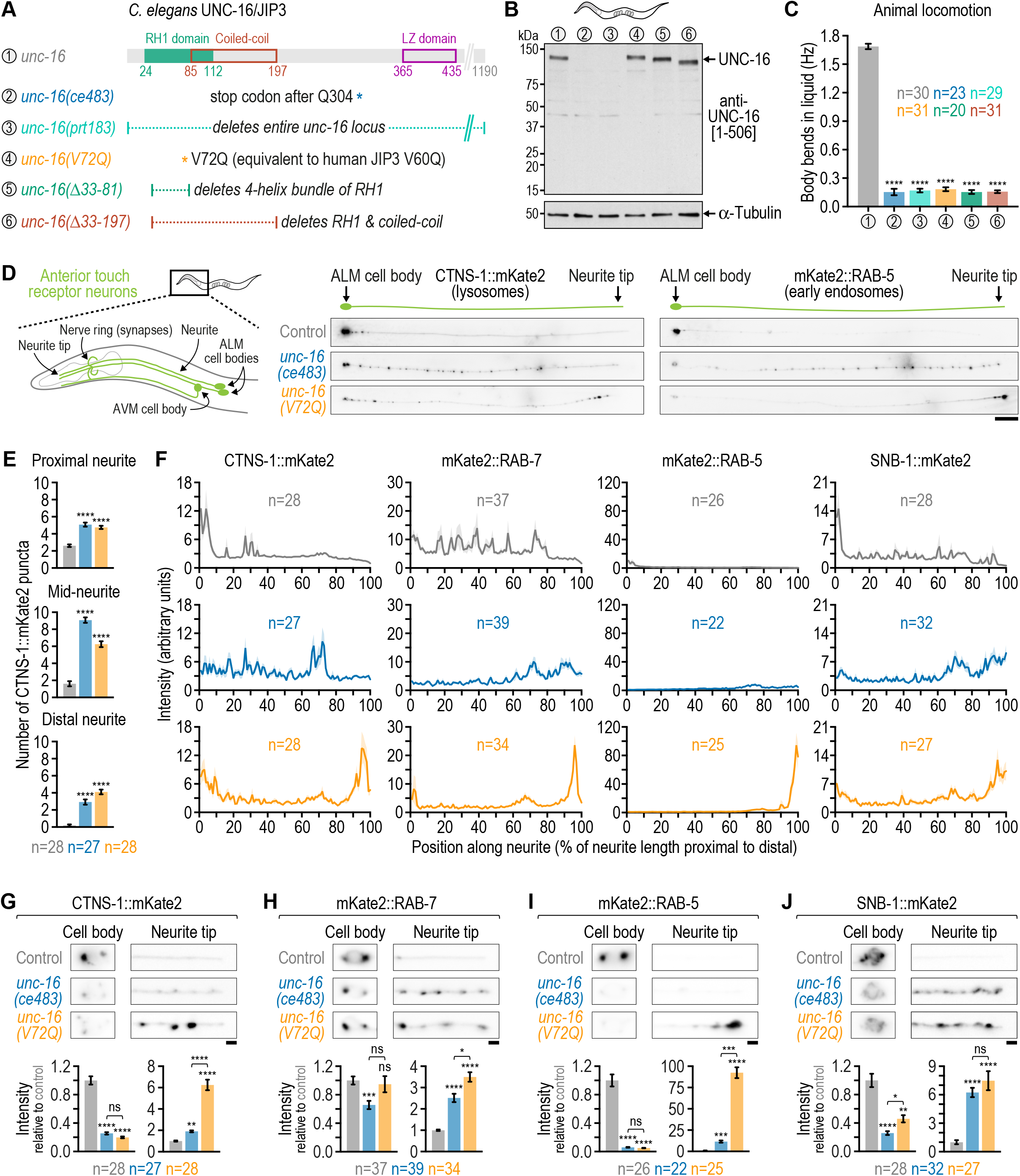
Displacing dynein light intermediate chain from *C. elegans* UNC-16/JIP3 through the UNC-16 V72Q mutation results in accumulation of endo-lysosomal organelles at the neurite tip of touch receptor neurons. **(A)** *(top)* Domain organization of the *C. elegans* UNC-16/JIP3 N-terminal region. Residue numbers correspond to isoform e (UniProt entry P34609-1). *(bottom)* Description of *unc: 16* mutants affecting the UNC-16 N-terminal region that are characterized in this study. **(B)** Immunoblot of adult *C. elegans* lysates using an affinity-purified rabbit polyclonal antibody raised against UNC-16 residues 1-506. The membrane was re-probed with an anti-α-tubulin antibody as a loading control. Molecular weight is in kilodaltons (kDa). **(C)** Locomotion of animals at the young adult stage, assessed by determining body bending frequency (mean ± SEM) in liquid medium. *n* denotes the number of animals examined. Statistical significance (wild-type N2 control versus *unc:16* mutants) was determined by ANOVA on ranks (Kruskal-Wallis nonparametric test) followed by Dunn’s multiple comparison test. \*\*\*\**P* < 0.0001. **(D)** *(left)* Location of the *C. elegans* anterior touch receptor neurons. ALM and AVM are the anterior lateral and anterior ventral mechanosensory neurons, respectively, which extend processes into the nose and the nerve ring. There are two ALM neurons, which are equivalent for the purpose of this study. Note that the neurite tip does not contain synapses, which are instead located in the nerve ring and were not imaged in this study. *(right)* Fluorescence images (maximum intensity z-stack projection, inverted grayscale) of the ALM neuron in L4 animals expressing a transgene-encoded marker for lysosomes (CTNS-1::mKate2) or early endosomes (mKate2::RAB-5) in touch receptor neurons. Scale bar, 20 µm. **(E)**Number of CTNS-1::mKate2 puncta (mean ± SEM) in the first quarter of ALM neurite length after the cell body (proximal neurite), the middle two quarters (mid-neurite), and the last quarter (distal neurite). *n* denotes the number of neurites examined (1 per animal). Statistical significance (control versus *unc:16* mutants) was determined as described for *(C)*. \*\*\*\**P* < 0.0001. **(F)** Fluorescence intensity profiles (mean ± SEM) along the ALM neurite in L4 animals expressing mKate2-tagged markers for endo-lysosomal organelles or synaptic vesicle precursors. *n* denotes the number of neurites examined (1 per animal). **(G) - (J)** *(top)* Fluorescence images (maximum intensity z-stack projection, inverted grayscale) of the ALM cell body and neurite tip. Scale bars, 2 µm. *(bottom)* Integrated fluorescence intensity (mean ± SEM; normalized to control) in the ALM cell body and the last 20 µm of the distal neurite (neurite tip). *n* denotes the number of neurites examined (1 per animal). Statistical significance was determined as described for *(C)*. \*\*\*\**P* < 0.0001; \*\*\**P* < 0.001; \*\**P* < 0.01; \**P* < 0.05; *ns* = not significant, *P* > 0.05.

### UNC-16(V72Q) causes prominent accumulation of endo-lysosomal organelles at the neurite tip of touch receptor neurons

To determine the consequences of the *unc:16(V72Q)* mutation for axonal transport, we set out to examine the distribution of organelles and vesicles in touch receptor neurons (TRNs). The 6 TRNs are situated underneath the cuticle and each have a long anteriorly directed neurite, in which microtubules are uniformly oriented toward the neurite tip (Arimoto *et al*., 2011). Therefore, kinesin-1 (along with other plus end-directed kinesins) and dynein mediate anterograde and retrograde transport in the neurite, respectively. We focused on the two ALM neurons, whose neurites extend into the head of the worm and branch off before the neurite tip to form synapses in the nerve ring (Fig. 4 D). To visualize different types of cargo, we used fluorescently tagged markers for lysosomes (CTNS-1::mKate2), late endosomes (mKate2::RAB-7), early endosomes (mKate2::RAB-5), early/recycling endosomes (mKate2::SYX-7), and synaptic vesicle precursors (SNB-1::mKate2), which were stably expressed in TRNs from the *mec:7* promoter after single-copy transgene integration using the MosSCI method (Frøkjær-Jensen *et al*., 2012). Consistent with prior work in motor neurons (Edwards *et al*., 2013), the *unc:16(ce483)* mutant increased the amount of CTNS-1::mKate2, mKate2::RAB-7, mKate2::RAB-5, and SNB-1::mKate2 in the ALM neurite at the larval L4 stage (Fig. 4 D; Fig. S4 A), which was quantified by counting the number of CTNS-1::mKate2 puncta present in the proximal, mid-, and distal neurite (Fig. 4 E); by determining the fluorescence intensity profile of the markers along the entire neurite (Fig. 4 F); and by measuring the integrated fluorescence intensity of the markers at the neurite tip and in the cell body (Fig. 4, G - J). In all cases, an increase of signal in the neurite was accompanied by a decrease of signal in the cell body.

We considered the possibility that *unc:16(ce483)* may not be a true null allele, since the nonsense mutation after Q304 and could in principle allow expression of an UNC-16 fragment that includes the RH1 domain and the first coiled-coil region. We therefore generated the knock-out allele *unc:16(prt183)* by removing the entire open reading frame and compared the intraneuronal distribution of mKate2::RAB-5 in *unc:16(prt183)* and *unc:16(ce483)* animals (Fig. S4 B). The two alleles had identical effects on mKate2::RAB-5 distribution, suggesting that *unc:16(ce483)* is essentially a null mutant. The identical appearance of *unc:16(ce483)* and *unc:16(prt183)* on immunoblots and their identical effect on animal locomotion further supports this idea (Fig. 4, B and C). Examination of mKate2::SYX-7 localization in the *unc: 16(prt183)* mutant showed that mKate2::SYX-7 puncta were broadly distributed throughout the ALM neurite, while mKate2::SYX-7 puncta in the control were almost exclusively found near the cell body (Fig. S5, B - D). Thus, the effect of *unc:16(prt183)* on mKate2::SYX-7 distribution is similar to that observed for other vesicle/organelle markers in the two *unc:16* null mutants. We have not examined mitochondria in *unc:16(ce483)* or *unc:16(prt183)* animals but note that increased mitochondrial density in TRN neurites has been reported for other *unc: 16* mutants (Sure *et al*., 2018).

Similar to the two *unc:16* null mutants, the *unc:16(V72Q)* mutant exhibited increased and decreased amounts of vesicle/organelle markers in the ALM neurite and cell body, respectively. However, with the exception of SNB-1::mKate2, the distribution profile of these markers in the neurite was distinct in the *unc:16(V72Q)* mutant. In contrast to *unc:16* null mutants, *unc:16(V72Q)* resulted in prominent accumulation of CTNS-1::mKate2, mKate2::RAB-7, mKate2::RAB-5, and mKate2::SYX-7 at the neurite tip (Fig. 4, D - J; and Fig. S5, B and D). The effect was particularly striking for mKate2::RAB-5. Thus, when compared to *unc:16* null mutants, *unc:16(V72Q)* shifts the distribution of endo-lysosomal markers in the neurite further in the anterograde direction.

The phenotype of the *unc:16(V72Q)* mutant is consistent with compromised retrograde transport. If this is indeed the case, dynein mutants should result in a similar accumulation of endo-lysosomal markers at the neurite tip. We therefore directly compared *unc:16(V72Q)* with *dli:1(F392A/F393A)*, which corresponds to the F447A/F448A mutation in human DLIC1 that abrogates binding to cargo adaptors (Celestino *et al*., 2019), including JIP3 as we show in this study (Fig. 2 G). We found that *unc:16(V72Q)* and *dli:1(F392A/F393A)* mutants have a similar distribution of CTNS-1::mKate2 and mKate2::RAB-5 along the neurite with a concomitant reduction of signal in the cell body (Fig. 5, A - C), although mKate2::RAB-5 accumulation at the neurite tip was not quite as pronounced in *dli:1(F392A/F393A)*. We note that since *dli:1(F392A/F393A)* animals are sterile, we could only analyze the first generation of homozygous mutants, which are likely to retain some residual maternal wild-type DLI-1. We conclude that mutating either the DLI-1 binding site in UNC-16 or the UNC-16 binding site in DLI-1 results in a similar distribution of endo-lysosomal organelles in TRNs.

**Figure 5.**
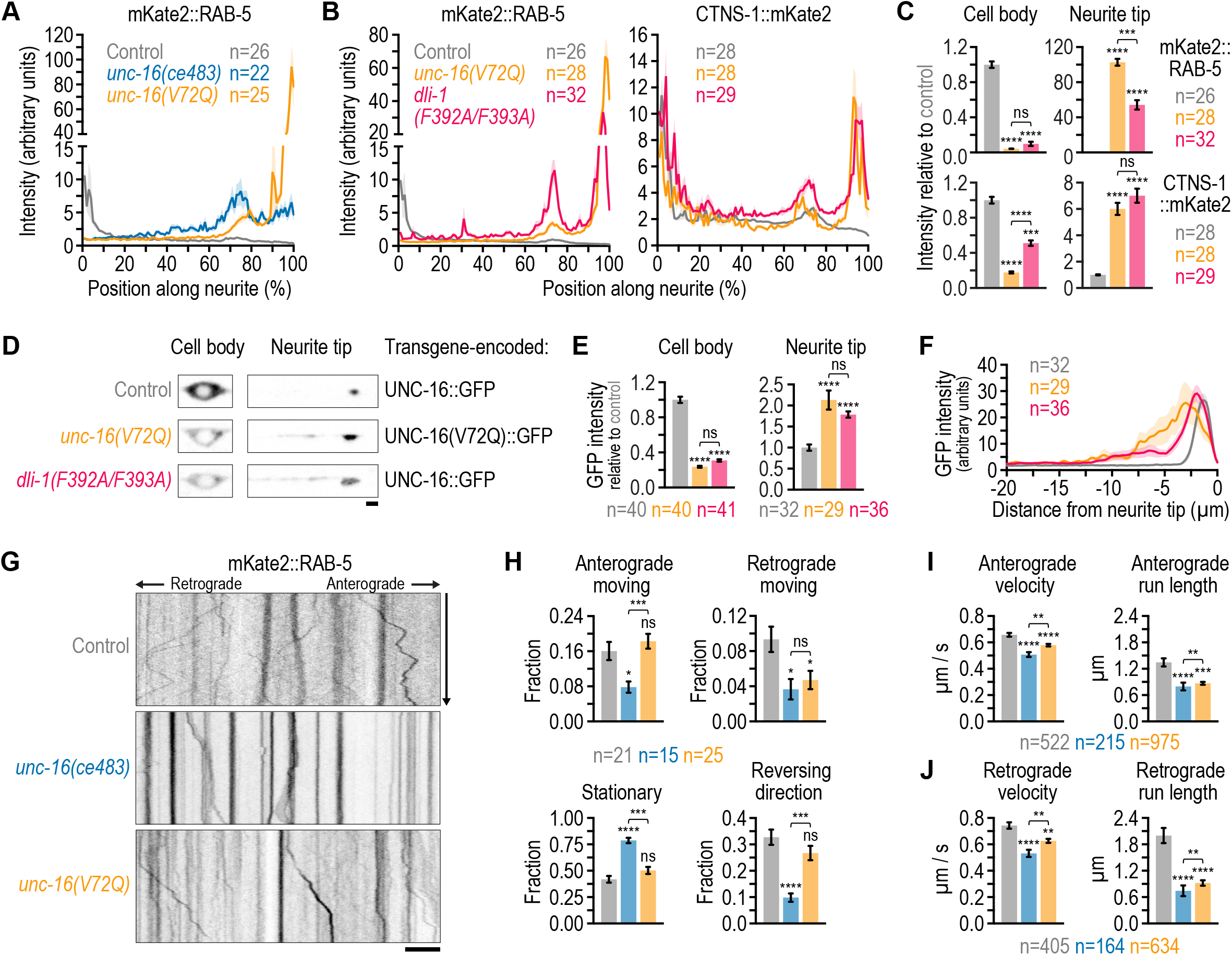
Mutations in the C-terminal helix 1 of dynein light intermediate chain and the UNC-16/JIP3 V72Q mutation result in similar transport defects in touch receptor neurons. **(A), (B)** Fluorescence intensity profiles (mean ± SEM) along the ALM neurite in L4 animals expressing a transgene-encoded marker for early endosomes (mKate2::RAB-5) or lysosomes (CTNS-1::mKate2). *n* denotes the number of neurites examined (1 per animal). Graph in *(A)* corresponds to the data in Fig. 4F, re-plotted with a split y-axis to highlight the difference between *unc:16(ce483)* and the control. **(C)** Integrated fluorescence intensity (mean ± SEM; normalized to control) in the ALM cell body and the last 20 µm of the distal neurite (neurite tip). *n* denotes the number of neurites examined (1 per animal). Statistical significance [control versus mutants and *unc:16(V72Q)* versus *dli:1(F392A/F393A)*] was determined by ANOVA on ranks (Kruskal-Wallis nonparametric test) followed by Dunn’s multiple comparison test. \*\*\*\**P* < 0.0001; \*\*\**P* < 0.001; *ns* = not significant, *P* > 0.05. **(D)** Fluorescence images (maximum intensity z-stack projection, inverted grayscale) of the ALM cell body and neurite tip in L4 animals expressing transgene-encoded UNC-16::GFP [background: endogenous wild-type *unc:16* with or without *dli:1(F392A/F393A)*] or UNC-16(V72Q)::GFP [background: endogenous *unc: 16(V72Q)*] in touch receptor neurons. Scale bar, 2 µm. **(E)** Integrated fluorescence intensity in the ALM cell body and neurite tip for the GFP-tagged UNC-16 versions described in *(D)*, plotted and statistically analyzed as in *(C)*. \*\*\*\**P* < 0.0001; *ns* = not significant, *P* > 0.05. **(F)** Fluorescence intensity profiles (mean ± SEM) at the ALM neurite tip for the GFP-tagged UNC-16 versions described in *(D)*. *n* denotes the number of neurites examined (1 per animal). **(G)** Fluorescence kymographs (inverted grayscale) of mKate2::RAB-5 particle motility in the ALM neurite, generated from time-lapse sequences (single z-section) recorded at the larval L2 stage. The imaged region is approximately 50 µm away from the cell body, which is located to the left. Scale bar, 5 µm. **(H) - (J)** Motility parameters (mean ± SEM) of mKate2::RAB-5 particles, derived from kymograph analysis. For *(H)*, *n* denotes the number of neurites examined (1 per animal). For *(I)* and *(J)*, *n* denotes the number of track segments, framed by a pause or a reversal, from at least 15 neurites (1 per animal). Statistical significance was determined as described for *(C)*. \*\*\*\**P* < 0.0001; \*\*\**P* < 0.001; \*\**P* < 0.01; \**P* < 0.05; *ns* = not significant, *P* > 0.05.

To examine how abrogating the interaction between DLI-1 and UNC-16 impacts the distribution of UNC-16 itself, we expressed transgenic UNC-16::GFP in TRNs from the *mec:4* promoter after single-copy integration. Consistent with previous reports on UNC-16::GFP localization (Byrd *et al*., 2001; Choudhary *et al*., 2017), UNC-16::GFP was detectable in the cell body (slightly enriched at what presumably is the golgi) and at the very tip of the neurite (Fig. 5, D - F). Expression of UNC-16(V72Q)::GFP in the *unc:16(V72Q)* background increased neurite tip levels, and a similar effect was observed for UNC-16::GFP in the *dli: 1(F392A/F393A)* background. This was accompanied by a decrease of signal in the cell body (Fig. 5, D - F). We conclude that abrogating the interaction between UNC-16 and DLI-1 increases the amount of UNC-16 in the neurite at the expense of cell body localization.

### The *unc/16(V72Q)* mutant exhibits anterograde bias of mKate2::RAB05 transport in TRN neurites

We next examined the transport kinetics of mKate2::RAB-5, which shows the most striking difference in distribution between *unc:16(V72Q)* and *unc:16* null mutants. Analysis of kymographs, constructed from time-lapse imaging sequences acquired in the middle segment of the ALM neurite at the larval L2 stage, showed that mKate2::RAB-5 particles exhibit bi-directional movement in control animals with ∼40 % of particles remaining stationary (Fig. 5, G and H). In the *unc:16(ce483)* mutant, the fraction of mKate2::RAB-5 particles moving exclusively in the anterograde or retrograde direction was decreased, and there were also less particles that reversed direction during the run. Consequently, more than 75 % of particles remained stationary for the duration of imaging (30 sec). The *unc:16(V72Q)* mutant showed a similar decrease in the fraction of exclusively retrograde moving particles but not in the fraction of exclusively anterograde moving particles, and there was no significant difference in the fraction of particles that reversed direction or remained stationary (Fig. 5, G and H). We also determined run length and velocity of mKate2::RAB-5 particles, both of which were decreased for either direction in the *unc:16(ce483)* and *unc:16(V72Q)* mutant (Fig. 5, I and J). We conclude that mKate2::RAB-5 particles are more mobile in the *unc:16(V72Q)* mutant than in the *unc:16(ce483)* mutant, and that transport in the *unc:16(V72Q)* mutant is biased in the anterograde direction relative to the control and the *unc:16(ce483)* mutant. This offers an explanation for the prominent accumulation of mKate2::RAB-5 that is observed at the neurite tip in the *unc:16(V72Q)* mutant.

### Accumulation of endo-lysosomal markers at the neurite tip in the *unc/16(V72Q)* mutant requires kinesin-1 activity

So far our analysis was consistent with the idea that displacing DLI-1 from UNC-16 in the *unc:16(V72Q)* mutant results in enhanced anterograde transport of endo-lysosomal organelles compared to what is observed in *unc:16* null mutants. To address whether kinesin-1 is implicated in this aspect of the *unc:16(V72Q)* phenotype, we used *unc:116(e2310)*, a reduction-of-function mutant of KHC (Patel *et al*., 1993). In *unc:16(V72Q)Runc:116(e2310)* double mutants, we no longer observed accumulation of CTNS-1::mKate2, mKate2::RAB-7, or mKate2::RAB-5 at the neurite tip, and (with the exception of mKate2::RAB-7) this was accompanied by an increase of signal in the cell body relative to the *unc:16(V72Q)* single mutant (Fig. 6, B and C). This shows that neurite tip accumulation of endo-lysosomal markers in the *unc:16(V72Q)* mutant requires kinesin-1 activity. Previous characterization of *unc: 116(e2310)* and other reduction-of-function alleles established that downregulation of kinesin-1 activity does not perturb microtubule orientation in the axon of DA9 (motor) and PHC (sensory) neurons (Yan *et al*., 2013), nor in the anterior neurite of ALM neurons (Arimoto *et al*., 2011). Thus, given that microtubule orientation likely remains correctly plus-end-out in the *unc:16(V72Q)Runc:116(e2310)* mutant, our result suggest that neurite tip accumulation of endo-lysosomal organelles in the *unc:16(V72Q)* mutant is the result of anterograde transport by kinesin-1.

**Figure 6.**
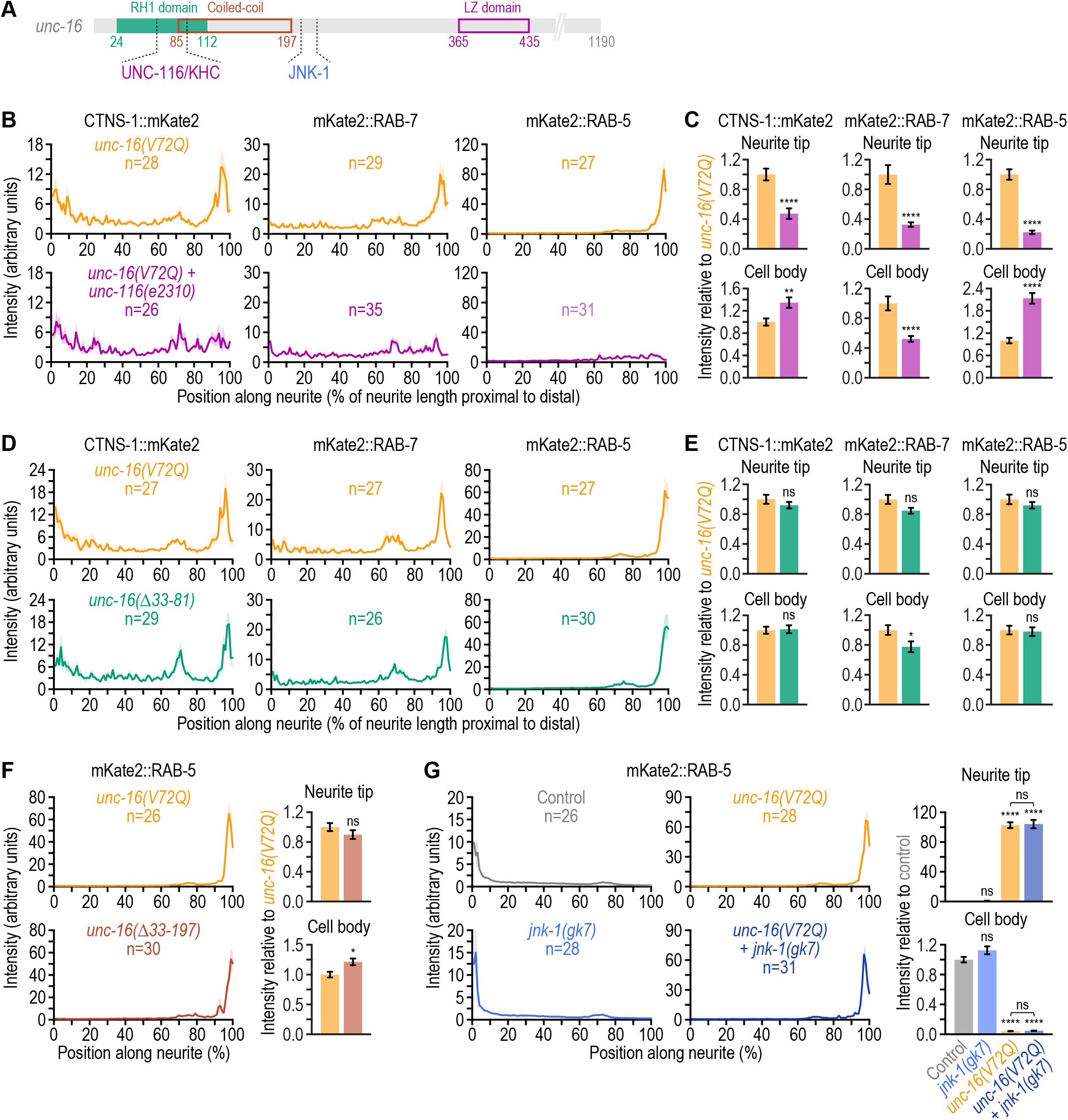
Neurite tip accumulation of endo-lysosomal organelles in the *unc/ 16(V72Q)* mutant requires kinesin-1 activity but not MAP kinase signalling, and N- terminal UNC-16 deletions mimic the *unc/16(V72Q)* mutant. **(A)** Domain organization of the N-terminal UNC-16 region with the putative binding sites for UNC-116/KHC and JNK-1. **(B), (D)** Fluorescence intensity profiles (mean ± SEM) along the ALM neurite in L4 animals expressing a transgene-encoded marker for lysosomes (CTNS-1::mKate2), late endosomes (mKate2::RAB-7), or early endosomes (mKate2::RAB-5) in touch receptor neurons. *n* denotes the number of neurites examined (1 per animal). **(C), (E)** Integrated fluorescence intensity [mean ± SEM; normalized to *unc:16(V72Q)*] in the ALM cell body and the last 20 µm of the distal neurite (neurite tip). The number of neurites examined in *(C)* and *(E)* corresponds to the number *n* in *(B)* and *(D)*, respectively. Statistical significance was determined by the Mann-Whitney test. \*\*\*\**P* < 0.0001; \*\**P* < 0.01; \**P* < 0.05; *ns* = not significant, *P* > 0.05. (F), (G) *(left)* Fluorescence intensity profiles (mean ± SEM) in the ALM neurite, as described for *(B)* and *(D)*. *(right)* Integrated fluorescence intensity [mean ± SEM; normalized to *unc:16(V72Q)* for *(F)* and to the control for *(G)*] in the cell body and at the neurite tip. The number of neurites examined corresponds to the number *n* for the fluorescence intensity profiles on the *(left)*. Statistical significance [control versus mutants and *unc:16(V72Q)* versus *unc: 16(V72Q)Rjnk:1(gk7)*] was determined by the Mann-Whitney test for *(F)* and by ANOVA on ranks (Kruskal-Wallis nonparametric test) followed by Dunn’s multiple comparison test for *(G)*. \*\*\*\**P* < 0.0001; \**P* < 0.05; *ns* = not significant, *P* > 0.05.

Neurite tip accumulation of endo-lysosomal markers could be driven by kinesin-1 that is directly bound to UNC-16. Alternatively, the relevant kinesin-1 pool could be associated with another adaptor that is (co-)dependent on UNC-16. Indeed, evidence from cultured mouse neurons suggests that JIP1 can bind to and cooperate with JIP3 in anterograde axonal transport (Sun *et al*., 2017; Hammond *et al*., 2008). To ask whether *C. elegans* JIP-1 is the source of the kinesin-1 activity that drives neurite tip accumulation of endo-lysosomal markers in the *unc:16(V72Q)* mutant, we generated the putative null allele *jip:1(prt187)* by deleting 7.2 kb of the open reading frame and introducing two nonsense mutations followed by a frameshift after residue L410, i.e. before the binding sites for KHC and KLC (Fig. S5 A). Analysis of intraneuronal mKate2::SYX-7 distribution showed that the *jip:1(prt187)* single and the *jip: 1(prt187)Runc:16(prt183)* double mutant were similar to the control and the *unc:16(prt183)* single mutant, respectively (Fig. S5, B - D). Although the neurite tip signal of mKate2::SYX-7 in the *jip:1(prt187)Runc:16(V72Q)* double mutant tended to be decreased relative to the neurite tip signal in the *unc:16(V72Q)* single mutant, the difference did not reach statistical significance. We conclude that JIP-1–bound kinesin-1 may make a modest contribution but is not the main driver of endo-lysosomal organelle accumulation at the neurite tip in the *unc: 16(V72Q)* mutant.

### Combining the UNC-16 V72Q mutation with a disease-associated mutation in UNC-16 that abrogates KLC binding recapitulates the *unc/16* null phenotype

Kinesin-1 interacts with JIP3/UNC-16 in two ways: through KHC, which binds to the highly conserved JIP3/UNC-16 RH1 domain (Sun *et al*., 2011 and this study), and KLC, which binds to the equally well conserved leucine zipper domain that precedes the RH2 domain (Fig. 1 A; Nguyen *et al*., 2005; Sakamoto *et al*., 2005). To examine the role of kinesin-1 that is bound to UNC-16, we first focused on the KHC interaction (Fig. 6 A). We reasoned that if the UNC-16 interaction with KHC is important for endo-lysosomal marker accumulation at the neurite tip in the *unc:16(V72Q)* mutant, this accumulation should not occur when the RH1 domain is deleted. We therefore generated animals expressing an UNC-16 mutant lacking residues 33-81, which deletes the two helices that form the 4-helix bundle in the RH1 dimer (Fig. 4 A). The deletion includes most of the region previously implicated in the interaction between KHC and mouse JIP3 (Sun *et al*., 2011; residues 50-80 in mouse correspond to residues 62-92 in *C. elegans*). Immunoblotting showed that UNC-16(Δ33-81) and wild-type UNC-16 are expressed at similar levels, and the body bending assay revealed severe locomotion deficiency in *unc:16(Δ33:81)* animals, which was indistinguishable from that of *unc:16(V72Q)* animals (Fig. 4, B and C). Analysis of CTNS-1::mKate2, mKate2::RAB-7, and mKate2::RAB-5 distribution in the ALM neuron showed that *unc:16(Δ33:81)* had the same effect as *unc:16(V72Q)* (Fig. 6, D and E). Since we could not rule out the possibility that UNC- 16(Δ33-81) has residual affinity for KHC, we also generated the mutant *unc:16(Δ33:197)*, which deletes the RH1 domain along with the entire first coiled-coil region of UNC-16 (Fig. 4 A). Immunoblotting showed that UNC-16(Δ33-197) levels were similar to those of wild-type UNC-16, and that *unc:16(Δ33:197)* animals were as locomotion-deficient as *unc:16(V72Q)* animals (Fig. 4, B and C). Strikingly, intraneuronal distribution of mKate2::RAB-5 in the *unc: 16(Δ33:197)* mutant was similar to that in the *unc:16(V72Q)* mutant (Fig. 6 F). This shows that simultaneous deletion of the binding sites for KHC and DLI-1 in UNC-16 mimics the specific loss of DLI-1 binding with regard to endo-lysosomal organelle distribution in TRNs.

Given that the predicted binding site for the MAP kinase JNK-1 is located just beyond the first coiled-coil region that is deleted in *unc:16(Δ33:197)*, we used the null mutant *jnk: 1(gk7)* to ask whether JNK signaling is involved in neurite tip accumulation of endo-lysosomal markers in the *unc:16(V72Q)* mutant (Villanueva *et al*., 2001). Analysis of intraneuronal mKate2::RAB-5 distribution showed that the *jnk:1(gk7)* single mutant is indistinguishable from the control, while the *jnk:1(gk7)*j*unc:16(V72Q)* double mutant is indistinguishable from the *unc:16(V72Q)* single mutant (Fig. 6 G). Thus, neurite tip accumulation of endo-lysosomal organelles in the *unc:16(V72Q)* mutant occurs independently of JNK-1 signalling.

The results with N-terminal deletion mutants of UNC-16 implied that the interaction between UNC-16(V72Q) and KHC is not required for endo-lysosomal organelle accumulation at the neurite tip. We therefore turned our attention to the KLC binding site in UNC-16 (Fig. 7 A). KLC binds via its TPR domain to the JIP3 leucine zipper, and this interaction has been analyzed at atomic resolution for mouse JIP3 (Cockburn *et al*., 2018). Recent work uncovered missense mutations in the human JIP3-encoding gene MAPK8IP that cause neurodevelopmental disorders and intellectual disability (Iwasawa *et al*., 2019; Platzer *et al*., 2019), and one of the mutations, L444P, is predicted to perturb KLC binding to JIP3 (Platzer *et al*., 2019). To test whether this is indeed the case, we generated purified recombinant human KLC1[185-501], which corresponds to the TPR domain, and mouse JIP3[186-505] with and without the L439P mutation, which is equivalent to human L444P (Fig. 7, A and B). Pull-downs with the GST-tagged KLC1 fragment demonstrated that mutating this highly conserved leucine to proline in JIP3 abrogates binding to KLC1 (Fig. 7 B).

**Figure 7.**
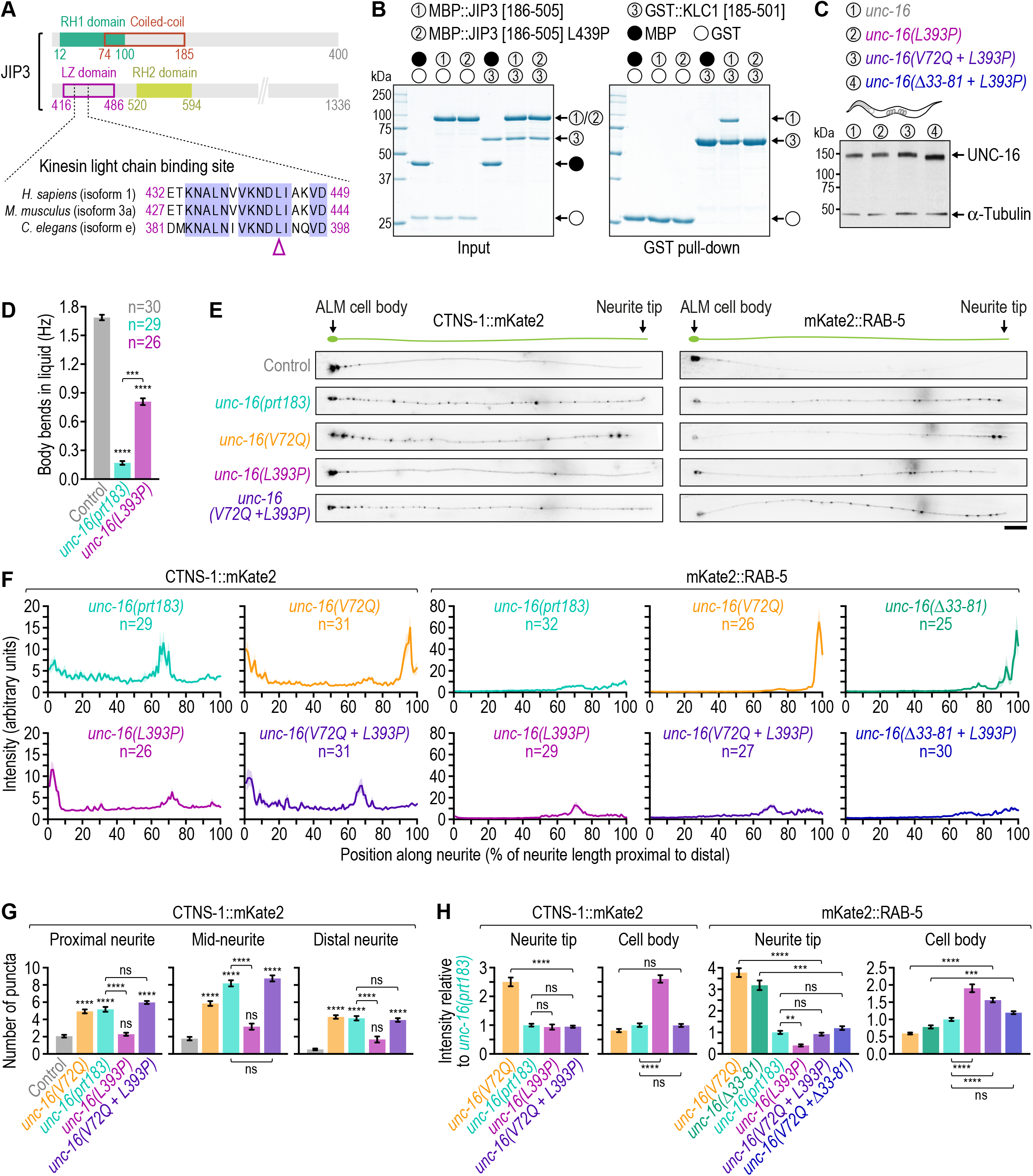
Neurite tip accumulation of endo-lysosomal organelles in the *unc/ 16(V72Q)* mutant requires the interaction between UNC-16/JIP3 and kinesin light chain. **(A)** Domain organization of human JIP3 with sequence alignment of the kinesin light chain binding site in the JIP3 leucine zipper (LZ) domain. Arrowhead points to the leucine residue in JIP3 (L444 in human; L439 in mouse; L393 in *C. elegans*) whose mutation to proline causes neurological disease and was predicted to interfere with kinesin light chain binding (Platzer *et al*., 2019). **(B)** *(left)* Coomassie Blue-stained SDS-PAGE gel of purified recombinant protein mixtures prior to addition of glutathione agarose resin (Input). *(right)* Coomassie Blue-stained SDS-PAGE gel of proteins eluted from glutathione agarose resin after GST pull-down. Proteins correspond to human kinesin light chain 1 (UniProt entry Q07866-1) and mouse JIP3 (Q9ESN9-5). Molecular weight is indicated in kilodaltons (kDa). **(C)** Immunoblot of *C. elegans* adult lysates using antibodies against UNC-16 and α-tubulin (loading control). Molecular weight is indicated in kilodaltons (kDa). **(D)** Locomotion of animals at the young adult stage, assessed by determining body bending frequency (mean ± SEM) in liquid medium. *n* denotes the number of animals examined. Statistical significance [wild-type N2 control versus *unc: 16* mutants and *unc:16(prt183)* versus *unc:16(L393P)*] was determined by ANOVA on ranks (Kruskal-Wallis nonparametric test) followed by Dunn’s multiple comparison test. \*\*\*\**P* < 0.0001; \*\*\**P* < 0.001. **(E)** Fluorescence images (maximum intensity z-stack projection, inverted grayscale) of the ALM neuron in L4 animals expressing a transgene-encoded marker for lysosomes (CTNS-1::mKate2) or early endosomes (mKate2::RAB- 5) in touch receptor neurons. Scale bar, 20 µm. **(F)** Fluorescence intensity profiles (mean ± SEM) along the ALM neurite in the animals described in *(E)*. *n* denotes the number of neurites examined (1 per animal). **(G)** Number of CTNS-1::mKate2 puncta (mean ± SEM) in the first quarter of ALM neurite length after the cell body (proximal neurite), the middle two quarters (mid-neurite), and the last quarter (distal neurite). The number of neurites examined (1 per animal) corresponds to the number *n* in *(F)*. Statistical significance (control versus *unc:16* mutants; other comparisons indicated by brackets) was determined by ANOVA on ranks (Kruskal-Wallis nonparametric test) followed by Dunn’s multiple comparison test. \*\*\*\**P* < 0.0001; *ns* = not significant, *P* > 0.05. **(H)** Integrated fluorescence intensity [mean ± SEM; normalized to *unc:16(prt183)*] in the ALM cell body and the last 20 µm of the distal neurite (neurite tip). The number of neurites examined corresponds to the number *n* in *(F)*. Statistical significance (control versus *unc:16* mutants; other comparisons indicated by brackets) was determined as described for *(G)*. \*\*\*\**P* < 0.0001; \*\*\**P* < 0.001; \*\**P* < 0.01; *ns* = not significant, *P* > 0.05.

In a previous effort to model the human disease, the analogous mutation (L393P) was introduced into *C. elegans* UNC-16, which resulted in an intermediate locomotion deficiency and a moderate increase in axonal lysosome density in cholinergic motor neurons (Platzer *et al*., 2019). We confirmed the intermediate locomotion deficiency of *unc:16(L393P)* animals (Fig. 7 D) and found that *unc:16(L393P)* increased CTNS-1::mKate2 and mKate2::RAB-5 levels in the ALM neurite, albeit to a lesser degree than in the null mutant *unc:16(prt183)* (Fig. 7, E - H). We conclude that the UNC-16(L393P) mutation produces a reduction-of-function phenotype, consistent with previous work (Platzer *et al*., 2019).

We then introduced the N-terminal mutations V72Q and Δ33-81 into the *unc: 16(L393P)* mutant background and confirmed by immunoblot that UNC-16(L393P) and the corresponding double mutant proteins were expressed at levels comparable to those of wild-type UNC-16 (Fig. 7 C). In contrast to the *unc:16(V72Q)* single mutant, the prominent neurite tip accumulation of CTNS-1::mKate2 and mKate2::RAB-5 was no longer observed in the double mutants, and the distribution in the ALM neuron now resembled that of the *unc:16* null mutant (Fig. 7, E - H). These results show that when the interaction between UNC-16 and DLI- 1 is perturbed, UNC-16 promotes neurite tip accumulation of endo-lysosomal organelles in a manner that depends on its association with KLC.

## DISCUSSION

In this study we identified the RH1 domain, whose presence defines a family of cargo adaptors for cytoskeletal motors, as novel binding partners for DLIC, and we elucidated key molecular determinants of the interaction. We then engineered a separation-of-function mutant of *C. elegans* UNC-16/JIP3 that cannot bind to DLIC, whose characterization revealed motor-dependent transport roles of JIP3 in neurons. For simplicity, we hereafter use the term JIP3 for both the vertebrate homolog and *C. elegans* UNC-16, unless we refer to a specific UNC- 16 mutation.

Prior work showed that the RH1 domain of JIP3 and RILPL2 binds to KHC and myosin Va, respectively (Sun *et al*., 2011; Wei *et al*., 2013), and we demonstrate here that the RH1 domain of human JIP3, JIP4, and RILP bind to DLIC. Thus, RH1 domains interact with all three types of cytoskeletal motors. We show that RILPL1’s N-terminal half also binds to DLIC, presumably via its RH1 domain, suggesting that RILPL1 is an adaptor for dynein. Based on structural work on the mouse RILPL2-myosin Va interaction (Wei *et al*., 2013), we mutated a conserved valine to glutamine in the RH1 domain of RILP (V59Q), JIP3 (V60Q), and JIP4 (V55Q). The point mutation abrogated the RH1–DLIC interaction, as did point mutations in helix 1 of the DLIC C-terminal tail. This suggests that the same hydrophobic pocket (one on each side of the RH1 4-helix bundle) that in RILPL2 accommodates helix 2 of the myosin Va globular tail domain is used in the dynein adaptors RILP, RILPL1, JIP3, and JIP4 to accommodate the amphipathic DLIC helix. Given the high sequence conservation between RH1 domains, this raises the question of how binding specificity for DLIC and myosin Va is achieved. The mouse RILPL2 study noted that F56, which participates in an additional hydrophobic interaction with myosin Va’s globular tail domain, is replaced by a proline in other members of the RH1 domain family (Wei *et al*., 2013). Interestingly, the F56P mutation abrogated RILPL2 binding to myosin Va, while the P55F mutation enabled mouse RILP to bind myosin Va with even higher affinity than RILPL2 (Wei *et al*., 2013). We find that the analogous P56F mutation in human RILP abrogates DLIC binding, consistent with its conversion to a myosin Va binder, but the converse is not the case: the F66P mutation is not sufficient to enable human RILPL2 to bind to DLIC (data not shown). This implies that there are additional distinguishing features in the RH1 domain of RILP and RILPL2 that confer binding specificity for DLIC and myosin Va, respectively. Unfortunately, we have so far failed to obtain crystals suitable for structural analysis of the RH1–DLIC interaction.

We show that JIP3’s RH1 domain not only binds DLIC but also the C-terminal tail domain of KHC, consistent with prior work (Sun *et al*., 2011), and that KHC binding is not affected by the JIP3 V60Q mutation. The residues in RH1 that mediate KHC binding remain to be identified, but the V60Q mutation and binding competition assays suggest that KHC and DLIC interact with distinct surfaces on the JIP3 RH1 domain. Simultaneous binding of KHC and DLIC to JIP3 could in principle permit kinesin-1 or dynein to remain associated with JIP3 while the other motor is actively engaged in transport. It is also tempting to speculate that the RH1 domain could play a role in coordinating the activities of the two opposite-polarity motors via long-range allostery.

The binding of DLIC and KHC to JIP3 is conserved in *C. elegans*, which enabled us to generate the separation-of-function mutant *unc:16(V72Q)* that specifically abrogates the interaction between UNC-16/JIP3 and DLI-1/DLIC. Characterization of this mutant in TRNs establishes that JIP3 binding to DLIC promotes the retrograde transport of lysosomes and endosomes and is therefore critical for JIP3’s previously described organelle clearance function (Edwards *et al*., 2013; Edwards *et al*., 2015). Additionally, our results suggest that JIP3–bound dynein promotes retrograde transport of synaptic vesicle precursors marked by SNB-1/synaptobrevin. SNB-1–marked vesicles are a well-established cargo of the kinesin-3 UNC-104, and the idea that JIP3–bound dynein opposes anterograde-directed motility of UNC-104 on synaptic vesicles fits well with the long-standing observation that SNB-1::GFP mis-localization in *unc:104* mutants can be partially suppressed by *unc:16* mutants (Byrd *et al*., 2001).

While both *unc:16* null mutants and the *unc:16(V72Q)* mutant have increased amounts of endo-lysosomal organelles in the anterior neurite of TRNs, organelle distribution within the neurite is strikingly different: abrogating the JIP3–DLIC interaction through the UNC-16(V72Q) mutation results in prominent organelle accumulation at the neurite tip, which is not observed in *unc:16* null mutants. Importantly, neurite tip accumulation can be rescued by the disease-associated JIP3 mutation L393P (human L444P), which we demonstrate abrogates the KLC- JIP3 interaction. So far, TRKB and JIP3 itself are the only firmly established cargos for JIP3–bound kinesin-1 (incidentally, TRKB is not conserved in *C. elegans*). Our results argue that JIP3–bound kinesin-1 also transports endo-lysosomal organelles and that this anterograde transport function is opposed by JIP3–bound dynein. We therefore propose that JIP3 uses the two opposite polarity motors to regulate organelle transport in a bi-directional manner. The contribution of JIP3–bound kinesin-1 to anterograde organelle transport presumably occurs against the backdrop of other as-yet-unidentified kinesin activity that acts on the same organelles and drives their accumulation in neurites when JIP3 is absent (Edwards *et al*., 2013; Edwards *et al*., 2015; Miller, 2017). We also note that while combining the V72Q mutation with the L393P mutation recapitulates the endo-lysosomal organelle distribution observed in *unc:16* null mutants, *unc:16(L393P)* on its own already causes some organelle accumulation in neurites. This is consistent with the previously described phenotype of this mutant in motor neuron axons (Platzer *et al*., 2019). A likely explanation is that kinesin-1 transports JIP3 into the neurite as cargo, and abrogating the JIP3–KLC interaction therefore decreases the amount of JIP3 that is available in the neurite to promote retrograde transport (Miller, 2017).

In contrast to what we observed for endo-lysosomal organelles, we found no obvious difference in the distribution of SNB-1::mKate2 between *unc:16* null mutants and the *unc: 16(V72Q)* mutant. This implies that in the case of this vesicular cargo JIP3–bound dynein is not opposed by JIP3–bound kinesin-1, which is consistent with UNC-104 being the kinesin responsible for anterograde SNB-1 transport. Whether or not JIP3–bound dynein and kinesin-1 act on the same type of membrane–bound cargo may in part be determined by the small GTPases that are thought to mediate JIP3 recruitment: for example, binding of Arf6 to JIP3 promotes the interaction with dynactin and is incompatible with KLC binding (Montagnac *et al*., 2009; Cockburn *et al*., 2018), which would preclude JIP3–bound kinesin-1 from acting on cargo to which JIP3 is recruited via Arf6. On the other hand, the binding of Rab10 or Rab36 to JIP3’s RH2 domain would in principle permit JIP3 to interact with either dynactin or KLC. Although JIP3 has been shown to associate with various membrane compartments by localization studies and biochemical fractionation, including lysosomes and early endosomes (Abe *et al*., 2009; Becker and Bonni, 2006; Cason *et al*., 2021; Cavalli *et al*., 2005; Choudhary *et al*., 2017; Drerup and Nechiporuk, 2013; Gowrishankar *et al*., 2017), the molecular pathways through which JIP3 is recruited to the organelles and vesicles whose transport it regulates remain to be defined.

Interestingly, we find that deleting JIP3’s RH1 domain and the adjacent coiled-coil region, which abrogates both DLIC and KHC binding, has the same effect on mKate2::RAB-5 distribution as UNC-16(V72Q). This implies that kinesin-1 binding to JIP3 via KLC is sufficient to drive endo-lysosomal organelle accumulation at the neurite tip in the absence of the JIP3–DLIC interaction. KHC binding to JIP3 has been implicated in activation of kinesin-1 motility (Sun *et al*., 2011; Watt *et al*., 2015), and it would therefore be interesting to examine organelle transport kinetics in a JIP3 mutant that cannot bind KHC but can still bind DLIC. Of note, our molecular characterization of the JIP3–DLIC interaction suggests that previously characterized N-terminal deletions in mouse JIP3 (Sun *et al*., 2011; Sun *et al*., 2013; Sato *et al*., 2015; Watt *et al*., 2015; Ma *et al*., 2017; Sun *et al*., 2017), which were presumed to specifically abrogate KHC binding, also abrogate DLIC binding.

The previous characterization of JIP3 as an activator of kinesin-1 motility provides an explanation for how JIP3’s interaction with KLC could promote the anterograde transport of endo-lysosomal organelles that we describe in this study (Sun *et al*., 2011; Watt *et al*., 2015). By contrast, how the JIP3–DLIC interaction promotes retrograde transport is less obvious. Activation of processive dynein motility occurs in the context of a tripartite complex consisting of the motor, dynactin, and a cargo adaptor. In all adaptors that have been demonstrated to be *bona fide* activators of dynein motility, the N-terminal coiled-coil region of the adaptor binds along the entire dynactin filament, which is 37 nm long. Adaptor binding to the dynactin filament is critical for formation of the dynein-dynactin-adaptor complex, because it stabilizes the interaction between the dynactin filament and the N-terminal tails of the dynein heavy chain dimer (Urnavicius *et al*., 2015). JIP3’s N-terminal coiled-coil is predicted to be far shorter than the dynactin filament (∼15 nm for mouse JIP3 [Vilela *et al*., 2019]), so structural considerations argue that JIP3 on its own is unlikely to be a potent activator of dynein motility. It will be important to clarify whether the previously described interaction between JIP3 and dynactin is direct (Cavalli *et al*., 2005; Montagnac *et al*., 2009), and, if it is, to directly test whether JIP3 binds dynactin in a manner that promotes formation of a motile dynein-dynactin-JIP3 complex. A likely possibility is that JIP3’s role in promoting retrograde transport involves cooperation with an activating adaptor, which could take one of two forms: JIP3 could aid in initial dynein-dynactin recruitment to cargo and then hand the (perhaps pre-activated) motor complex over to an activating adaptor, or JIP3 could incorporate into the dynein-dynactin transport machine together with an activating adaptor. In the latter scenario, the additional binding sites for DLIC provided by the JIP3 RH1 domain could help recruit a second dynein to dynactin for increased force production and speed (Urnavicius *et al*., 2018). Elucidating the functional relationship between JIP3 and other dynein adaptors for endo-lysosomal organelles (including RILP, which, interestingly, suffers from the same structural shortcoming as JIP3 with regard to its potential as an activating adaptor) is an important future research direction.

## MATERIALS AND METHODS

### C. elegans strains

Worm strains (Table S1) were maintained at 20 °C on standard NGM plates seeded with OP50 *Escherichia coli* bacteria. The mutants *unc:16(prt183)*, *unc:16*(*V72Q)*, *unc:16(Δ33: 81)*, *unc:16(Δ33:197)*, and *jip:1(prt187)* were generated by CRISPR/Cas9-mediated genome editing, as described previously (Arribere *et al*., 2014; Paix *et al*., 2014). Genomic sequences targeted by sgRNAs are listed in Table S2. The modifications were confirmed by sequencing and strains were outcrossed 4 - 6 times against the wild-type N2 strain. Fluorescent markers were subsequently introduced by mating.

### Image acquisition

#### Animal locomotion assay

L4 hermaphrodites were transferred to a new NGM plate with bacteria 24 h before performing the assay. For imaging, animals were transferred to a slide containing a 2-µL drop of M9. Movements were tracked at 20 °C for 1 min at 40 frames per second using a SMZ 745T stereoscope (Nikon) with a QIClic CCD camera (QImaging) controlled by Micro-Manager software (Open Imaging). The wrMTrck plugin for Image J was used for automated counting of body bends.

#### Touch receptor neurons

To image mKate2–tagged organelle and vesicle markers in the ALM neuron, L4 hermaphrodites were paralyzed in M9 buffer containing 50 mM NaN_3_ for 5 min, transferred to a freshly prepared 2 % (w/v) agarose pad, and covered with an 18 mm x 18 mm coverslip (No. 1.5H, Marienfeld). Imaging was performed on an Axio Observer microscope (Zeiss) equipped with an Orca Flash 4.0 camera (Hamamatsu) and an HXP 200C Illuminator (Zeiss). A z-stack (step size 0.5 µm) that captured the entire neuron was acquired at 1 x 1 binning with a 40x NA 1.3 Plan-Neofluar objective. Image acquisition was controlled by ZEN 2.3 software (Zeiss). For time-lapse imaging of mKate2::RAB-5 particles in the ALM neuron, L2 hermaphrodites were paralyzed with 5 mM levamisole in M9 buffer for 10 min and mounted on agarose pads as described above. Only morphologically healthy neurites were imaged. Time-lapse sequences of a region approximately 50 µm away from the cell body were recorded in a temperature-controlled room at 20 °C on a Nikon Eclipse Ti microscope coupled to an Andor Revolution XD spinning disk confocal system, composed of an iXon Ultra 897 CCD camera (Andor Technology), a solid-state laser combiner (ALC-UVP 350i, Andor Technology), and a CSU-X1 confocal scanner (Yokogawa Electric Corporation). The system was controlled by Andor IQ3 software (Andor Technology). A single image was acquired every 200 ms for a total of 30 s at 1 x 1 binning using a 100x NA 1.45 Plan-Apochromat objective.

To image GFP–tagged UNC-16 in the ALM neuron, L4 hermaphrodites were paralyzed with NaN_3_ as described above and imaged on the spinning disc confocal microscope. Z-stacks (step size 0.2 µm) that captured the neurite tip and cell body were acquired at 1 x 1 binning with a 60x NA 1.4 Plan-Apochromat objective.

### Image analysis

Image analysis was performed using Fiji software (Image J version 1.52d).

#### Fluorescence intensity profile along the neurite

A segmented line with a width of 10 pixels was traced on top of the ALM neurite and the fluorescence intensity profile of the entire neurite was recorded using Fiji’s “Plot Profile” function. A parallel segmented line was traced adjacent to the neurite to obtain a background intensity measurement for each position along the neurite. The final fluorescence intensity at each position was calculated by subtracting the background signal from the neurite signal. Absolute positions along the neurite were normalized to neurite length with - % corresponding to the proximal neurite at (but not including) the cell body and 100 % corresponding to the distal neurite tip. The signal was averaged over 1 % intervals.

#### Integrated fluorescence intensity at neurite tip and in cell body

The integrated fluorescence intensity at the ALM neurite tip was determined for the last 20 µm of the neurite by summing up the fluorescence intensity in the profile plot described above and by subtracting the corresponding background signal. For mKate2::SYX-7, a different approach was necessary, since the diffuse signal in the neurite was too dim to accurately trace the neurite for profile plots. The integrated fluorescence intensity at the neurite tip, which was not visible in the control, was therefore measured in a 185 x 15-pixel (30 x 2.5 µm) ROI placed adjacent to the pharyngeal procorpus identified in the differential interference contrast (DIC) channel. The ROI was expanded by a few pixels on each side, and the difference in signal between the outer and inner ROI was used to define the integrated background intensity after normalization to the area of the inner ROI. The final integrated intensity was calculated by subtracting the integrated background intensity from the integrated intensity of the inner ROI. An analogous approach was used for the cell body with the inner ROI corresponding to the cell body outline.

#### Counting of CTNS-1::mKate2 and mKate2::SYX-7 puncta

For CTNS-1::mKate2 an empirically determined threshold (approximately 1.5-fold above the local diffuse background signal) was applied to the fluorescence intensity profile plot to determine the number and position of puncta, which were then grouped according to their position along the neurite: proximal neurite (≤ 25 % length), mid-neurite (> 25 % to ≤ 75 % length), and distal neurite (> 75 % length). Since we could not generate fluorescence intensity profile plots in neurites expressing mKate2::SYX-7, mKate2::SYX-7 puncta were counted directly in equally scaled fluorescence images, and, to define neurite length in the control, the neurite tip was estimated to lie at the transition between buccal cavity and pharyngeal procorpus identified in the DIC channel.

#### Motility parameters of mKate2::RAB-5 particles

Kymograph generation and analysis were performed with KymoAnalyzer (Neumann *et al*., 2017). Only time-lapse sequences during which there was no discernable movement of the animal and which had uniformly bright mKate2 signal along the neurite were considered. A track is defined as a single particle trajectory, and a segment corresponds to a portion within the track of a moving particle that is framed by a pause or a reversal. Values for run length and velocity reported in this study are for segments. Particles classified as “anterograde moving” or “retrograde moving” did so exclusively, i.e. their track did not contain any segments in which movement was reversed. Particles that had both anterograde and retrograde segments in their track were classified as “reversing direction”. Particles were classified as “stationary” if their track did not contain any moving segments.

### Antibody against UNC-16

Polyclonal antibodies against an N-terminal fragment of UNC-16 (residues 1-506) were generated as follows: GST::UNC-16(1-506) was expressed from pGEX6P-1 in bacteria, purified as described below, and injected into rabbits (GeneCust). For affinity purification of the antibodies, the GST fusion protein was cleaved using Prescission protease, and UNC- 16(1-506) was covalently coupled to a 1-mL HiTrap *N*-hydroxysuccinimide column (GE Healthcare).

### Immunoblotting

For immunoblots of *C. elegans* lysate, 100 adult hermaphrodites were collected into 1 mL of M9 buffer and washed with 3 x 1 mL M9 buffer and 3 x 1 mL M9 buffer containing 0.05 % Triton X-100. To 100 µL of worm suspension, 33 µL 4x SDS PAGE sample buffer [250 mM Tris-HCl pH 6.8, 30 % (v/v) glycerol, 8 % (w/v) SDS, 200 mM DTT, 0.04 % (w/v) bromophenol blue] and 20 µL of glass beads were added. Samples were incubated for 3 min at 95 °C and vortexed for 5 min with intermittent heating. After centrifugation at 20’000 x g for 1 min at room temperature, proteins in the supernatant were resolved on a 4 - 20 % gradient gel (Bio-Rad). Proteins were transferred to a 0.2-µm nitrocellulose membrane (GE Healthcare). The membrane was blocked with 5 % (w/v) non-fat dry milk in TBST (20 mM Tris-Cl, 140 mM NaCl, 0.1 % Tween, pH 7.6) and incubated overnight at 4 °C in TBST/5 % dry milk with rabbit polyclonal anti-UNC-16 antibody GC23 (1:1’200; made in-house), mouse monoclonal anti-α-tubulin antibody B512 (1:5’000; Sigma-Aldrich), rabbit polyclonal anti-GST antibody GC3 (1:7’500; Gama *et al*., 2017), or mouse monoclonal anti-6xHis antibody His.H8 (1:2’500; Millipore). The membrane was rinsed 5 x with TBST and incubated in TBST/5 % dry milk with goat secondary antibodies coupled to HRP (1:10’000; Jackson ImmunoResearch) for 1 h at room temperature. After 3 washes with TBST, proteins were visualized by chemiluminescence using Pierce ECL Western Blotting Substrate (Thermo Fisher Scientific) and x-ray film (GE Healthcare). Immunoblotting with the mouse monoclonal anti-Strep-tag II antibody StrepMAB (1:1’000; IBA) was carried out as described previously (Celestino *et al*., 2019).

### Plasmids for recombinant protein expression

cDNA encoding for the protein fragment to be expressed was inserted into a 2CT vector [N-terminal 6xHis::maltose binding protein (MBP) followed by a TEV protease cleavage site and C-terminal Strep-tag II] or into pGEX-6P-1 [N-terminal glutathione S-transferase (GST) followed by a Prescission protease cleavage site and C-terminal 6xHis]. Residue numbers in text and figures correspond to the following UniProt entries: Q96NA2-1 (human RILP), Q9JJC6-1 (mouse RILPL1), Q969X0-1 (human RILPL2), Q9UPT6-1 (human JIP3), Q9ESN9-5 (mouse JIP3), O60271-1 (human JIP4), Q9Y6G9-1 (human DYNC1LI1), Q07866- 1 (human KLC1), P28738-1 (mouse KIF5C), P34609-1 (*C. elegans* UNC-16), P34540-1 (*C. elegans* UNC-116), G5ED34-1 (*C. elegans* DLI-1).

### Recombinant protein expression and purification

Expression vectors were transformed into *E. coli* strains BL21, BL21-CodonPlus-RIL, or Rosetta. Expression was induced at an OD_600_ of 0.9 with 0.1 mM IPTG. After expression overnight at 18 °C, cells were harvested by centrifugation at 4’000 x g for 20 min.

For purification of proteins expressed from the 2CT vector, bacterial pellets were resuspended in lysis buffer A (50 mM HEPES, 250 mM NaCl, 10 mM imidazole, 1 mM DTT, 1 mM PMSF, 2 mM benzamidine-HCl, pH 8.0), lysed with a cell cracker, and cleared by centrifugation at 40’000 x g for 45 min at 4°C. Proteins were purified by tandem affinity chromatography using HisPur Ni-NTA resin (Thermo Fisher Scientific) followed by strep-tactin sepharose resin (IBA). Ni-NTA resin was incubated in batch with the cleared lysate in the presence of 0.1 % (v/v) Tween 20 for 1 h at 4° C and washed with wash buffer A (25 mM HEPES, 250 mM NaCl, 25 mM imidazole, 0.1 % Tween 20, 1 mM DTT, 2 mM benzamidine-HCl, pH 8.0). Proteins were eluted on a gravity column with elution buffer A (50 mM HEPES, 150 mM NaCl, 250 mM imidazole, 1 mM DTT, 2 mM benzamidine-HCl, pH 8.0). Fractions containing the recombinant protein were pooled, incubated overnight at 4°C with TEV protease (if 6xHis::MBP needed to be cleaved off), incubated in batch with strep-tactin sepharose resin for 1 h at 4° C, and washed with wash buffer B (25 mM HEPES, 250 mM NaCl, 0.1 % Tween 20, 1 mM DTT, 2 mM benzamidine-HCl, pH 8.0). Proteins were eluted on a gravity column with elution buffer B (100 mM Tris-HCl, 150 mM NaCl, 1 mM EDTA, 10 mM desthiobiotin, pH 8.0). Some proteins were further purified by size-exclusion chromatography on a Superose 6 10/300 or Superdex 200 Increase column (GE Healthcare) equilibrated with storage buffer (25 mM HEPES, 150 mM NaCl, pH 7.5) or ITC buffer (25 mM HEPES, 100 mM KCl, 0.5 mM TCEP, pH 7.5). Alternatively, the eluate from the gravity column was directly dialyzed against storage buffer. Glycerol and DTT were added to a final concentration of 10 % (v/v) and 1 mM, respectively (except for ITC samples), and aliquots were flash frozen in liquid nitrogen and stored at −80°C.

For purification of proteins expressed from the pGEX-6P-1 vector, bacterial pellets were resuspended in lysis buffer B (50 mM HEPES, 250 mM NaCl, 10 mM EDTA, 10 mM EGTA, 1 mM DTT, 1 mM PMSF, 2 mM benzamidine-HCl, pH 8.0), lysed with a cell cracker, and cleared by centrifugation at 40’000 x g for 45 min at 4°C. Proteins were purified by tandem affinity chromatography using glutathione agarose resin (Thermo Fisher Scientific) followed by Ni-NTA resin. Glutathione agarose resin was incubated in batch with the cleared lysate in the presence of 0.1 % (v/v) Tween 20 for 1 h at 4° C, washed with wash buffer C (25 mM HEPES, 250 mM NaCl, 0.1 % Tween 20, 1 mM DTT, 2 mM benzamidine-HCl, pH 8.0), and proteins were eluted on a gravity column with elution buffer C (50 mM HEPES, 150 mM NaCl, 10 mM reduced L-glutathione, 1 mM DTT, 2 mM benzamidine-HCl, pH 8.0). Alternatively, the resin was incubated overnight at 4°C with Prescission Protease in cleavage buffer (25 mM HEPES, 150 mM NaCl, 0.01 % Tween 20, 1 mM DTT, 2 mM benzamidine-HCl, pH 7.6) to remove the GST tag. Fractions containing the (cleaved) recombinant protein were pooled, incubated in batch with Ni-NTA resin for 1 h at 4° C, washed with wash buffer A, and eluted on a gravity column with elution buffer A. Proteins were further purified by size-exclusion chromatography or dialyzed before flash freezing, as described above.

### Size exclusion chromatography

For binding assays, proteins mixtures were incubated on ice for 30 min prior to size exclusion chromatography, which was performed at room temperature on an ÄKTA Pure 25L1 system with a Superdex 200 Increase column (GE Healthcare). Elution of proteins was monitored at 280 nm. 20 µL of successive 0.5-mL elution fractions were separated by SDS- PAGE and proteins were visualized by Coomassie Blue staining.

### Pull-down assays

Purified recombinant proteins were mixed in 45 µL pull-down buffer (50 mM HEPES, 100 mM NaCl, 5 mM DTT, pH 7.5). This initial mix was split into two halves, one corresponding to the input and the other to the pull-down. For pull-downs, the protein mix was incubated for 1 h at 4° C in 150 µL pull-down buffer supplemented with 15 µL glutathione agarose resin for GST pull-downs or amylose resin (New England Biolabs) for MBP pull-downs. After washing the resin with 3 x 500 µL pull-down buffer, proteins were eluted with the same buffer supplemented with 15 mM reduced L-glutathione (glutathione agarose resin) or 10 mM maltose (amylose resin). Eluted proteins were separated by SDS-PAGE gel and visualized by Coomassie Blue staining or immunoblotting.

### Isothermal titration calorimetry

ITC data was recorded in a MicroCal VP-ITC calorimeter (Malvern). All proteins were purified in the same batch of ITC buffer (25 mM HEPES, 100 mM KCl, 0.5 mM TCEP, pH 7.5). Titrations were performed at 25 °C and consisted of an initial 2 µL injection followed by a series of 28 injections of 10 µL each. In the cases where an interaction was detected a blank run was performed by titrating the titrant into ITC buffer. Data were processed with the ORIGIN software package (OriginLab) and in cases where an interaction was detected either the blank run was subtracted from the initial isotherm (to correct for the heat of dilution) or data were further processed for global fitting with the NITPIC, SEDPHAT, and GUSSI software packages.

### Analytical ultracentrifugation (AUC)

Sedimentation velocity AUC was performed at 42’000 rpm at 20 °C in an An-60 Ti rotor (Beckman Coulter) using standard double-sector centerpieces. Protein samples were diluted in AUC buffer (50 mM HEPES, 100 mM NaCl, 0.5 mM TCEP, pH 7.5). Approximately 300 radial absorbance scans at 280 nm were collected per each run with a time interval of 1 min. Analysis was performed using the SEDFIT suite to obtain the continuous distribution function of sedimentation coefficients [c(S)]. The software SEDNTERP was used to estimate buffer density and viscosity, as well as the protein partial specific volume. GUSSI software was used to generate the final figures.

### Statistical analysis

Statistical analysis was performed with Prism 8.0 software (GraphPad). The analytical method used is specified in the figure legends.

## ACKNOWLEDGEMENTS

The authors thank Stacey L. Edwards and Kenneth G. Miller (Oklahoma Medical Research Foundation) for *C. elegans* strain KG5309. Some strains were provided by the CGC, which is funded by NIH Office of Research Infrastructure Programs (P40 OD010440).

---

**Figure S1.**
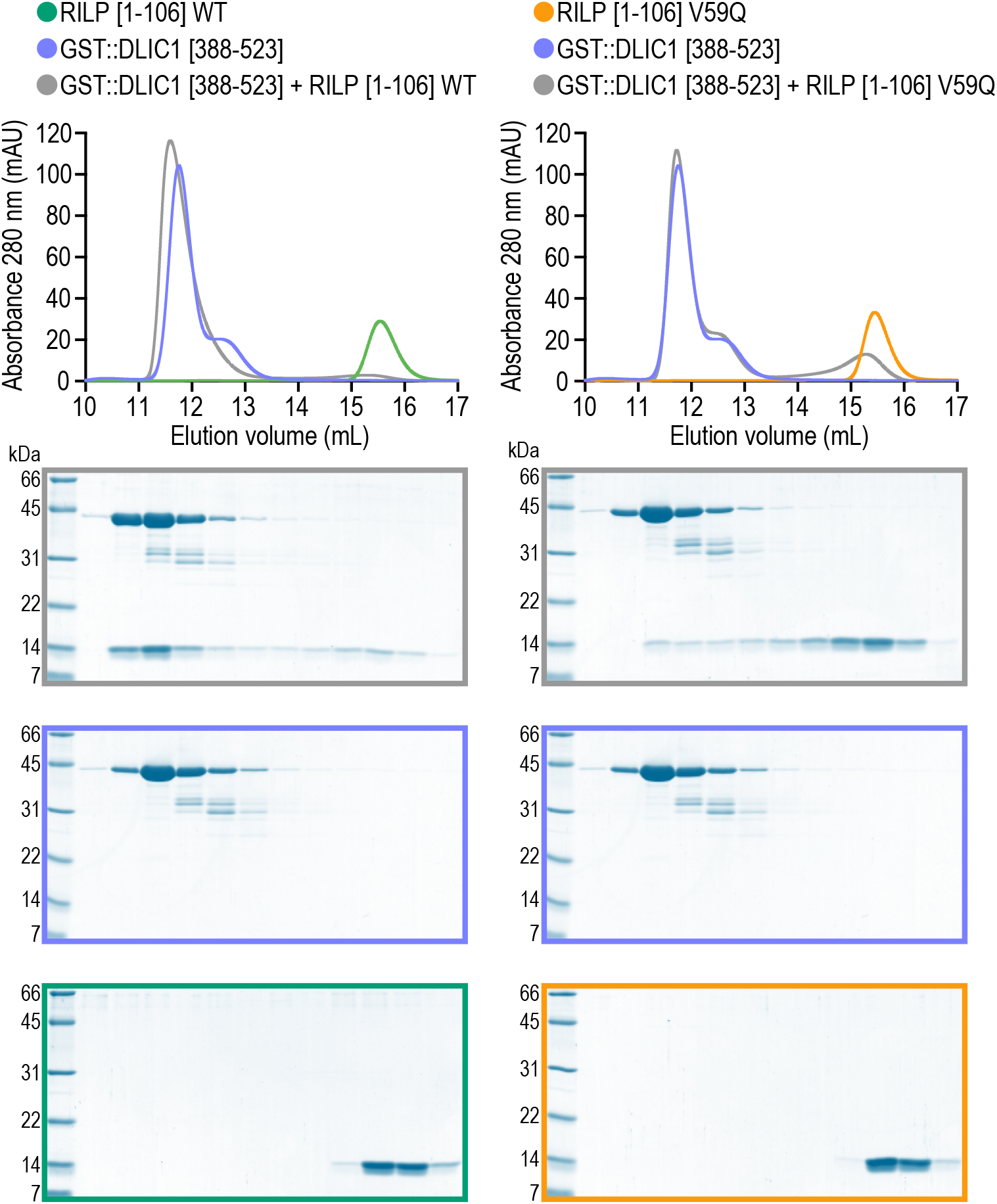
The V59Q mutation in RILP reduces the affinity for dynein light intermediate chain. Elution profiles *(top)* and Coomassie Blue-stained SDS-PAGE gels *(bottom)* of purified recombinant proteins after size exclusion chromatography on a Superdex 200 Increase column. Proteins correspond to the human homologs described in Fig. 1A. WT denotes wild-type. Molecular weight is indicated in kilodaltons (kDa).

**Figure S2.**
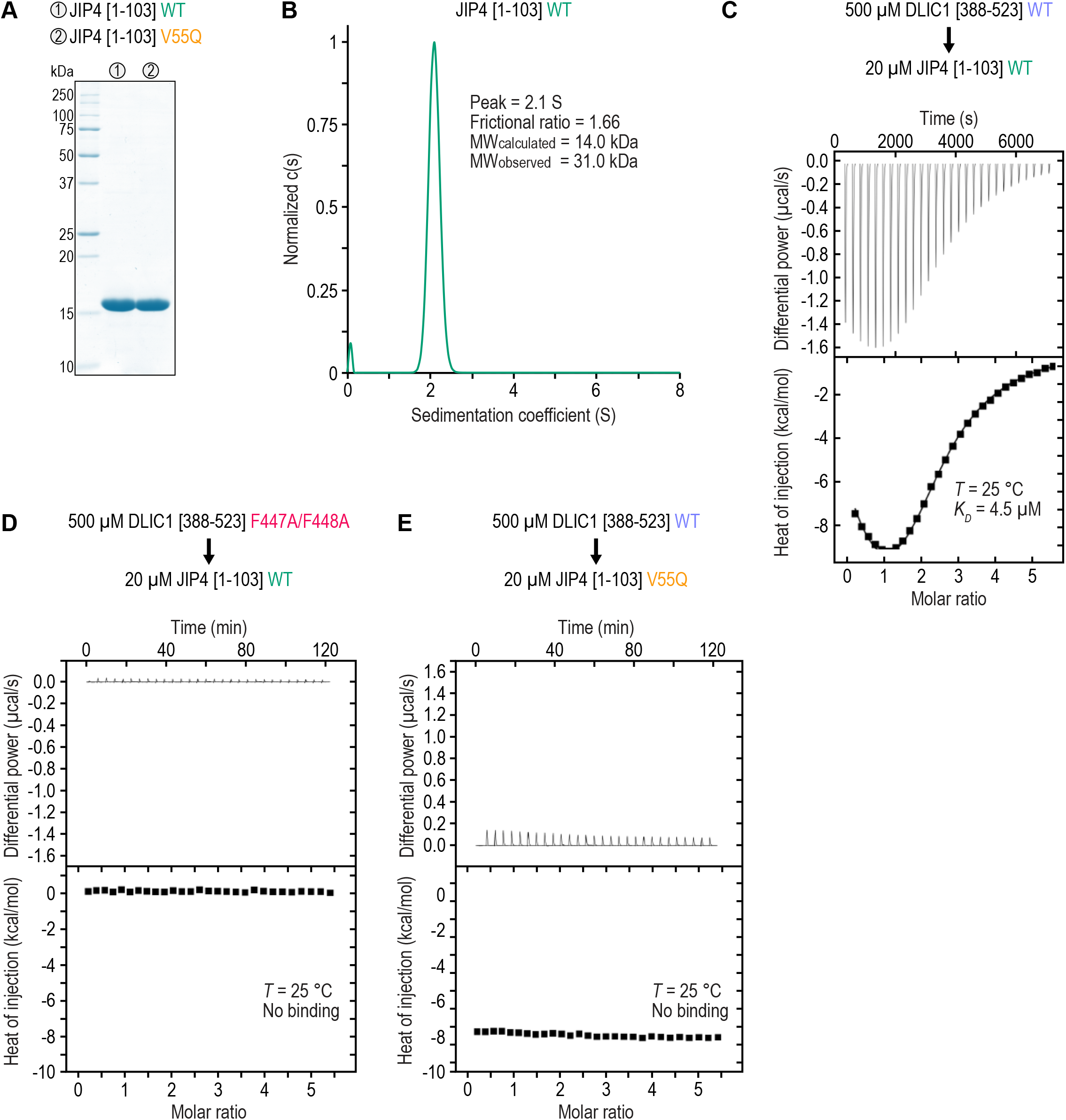
The V55Q mutation in JIP4 abrogates the binding to dynein light intermediate chain. **(A)** Coomassie Blue-stained SDS-PAGE gel of purified recombinant proteins used in AUC and ITC experiments. Proteins correspond to the human homologs described in Fig. 1A. WT denotes wild-type. Molecular weight is indicated in kilodaltons (kDa). **(B)** Sedimentation velocity AUC profile with theoretical (MW_calculated_) and experimentally measured molecular mass (MW_observed_). The MW_observed_ value indicates that the protein is dimeric in solution. **(C) - (E)** Thermograms and binding isotherms of representative ITC titrations. JIP4[1-103] concentration is the concentration of the dimer. The data in *(C)* were best described by a model corresponding to two symmetric sites with a single macroscopic dissociation constant (*K_D_*) using the SEDPHAT software package. The *K_D_* value was determined by global fitting of two independent runs with a 68.3 % confidence interval of 2.8 - 7.4 µM.

**Figure S3.**
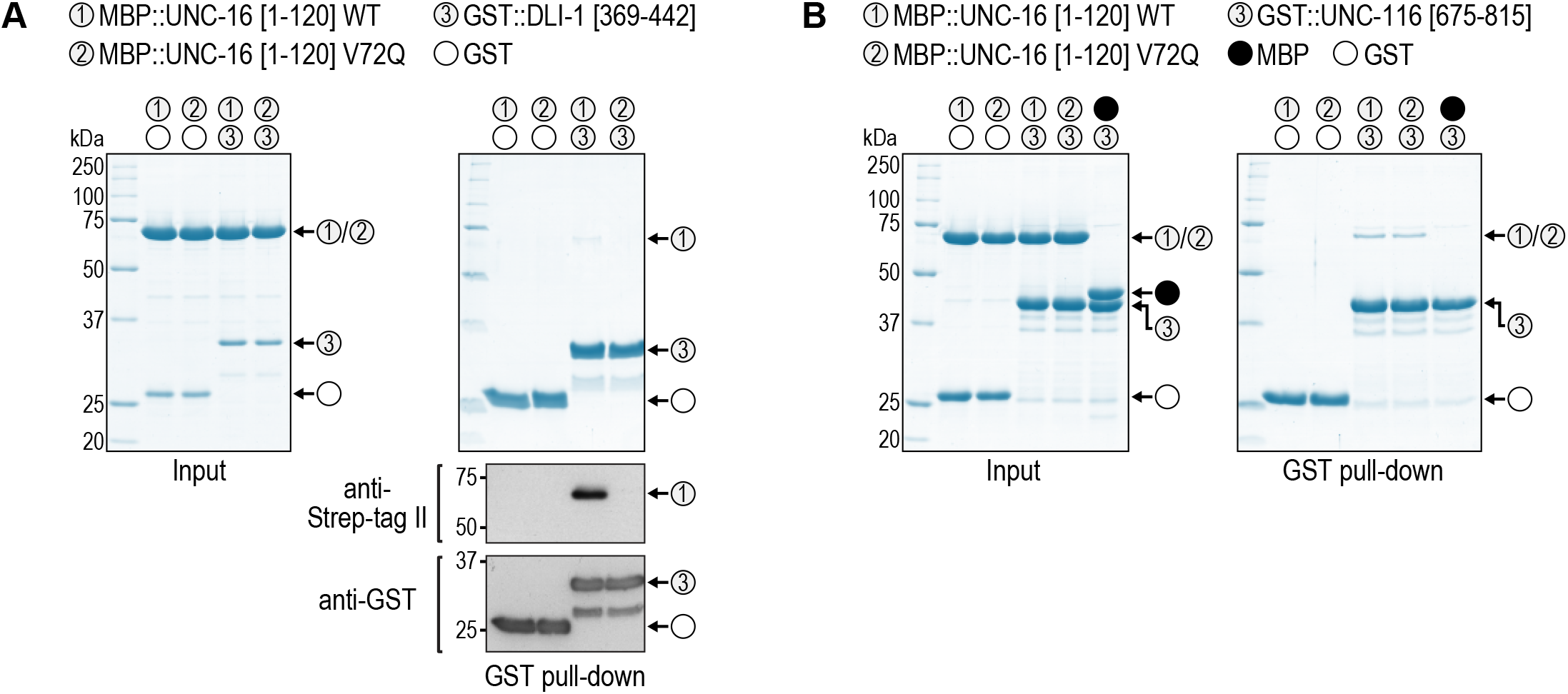
Human JIP3 V60Q is equivalent to *C. elegans* UNC-16 V72Q. **(A)**, **(B)** *(left)* Coomassie Blue-stained SDS-PAGE gel of purified recombinant protein mixtures prior to addition of glutathione agarose resin (Input). *(right)* Coomassie Blue-stained SDS-PAGE gel of proteins eluted from glutathione agarose resin after GST pull-down. Proteins correspond to the *C. elegans* homologs of JIP3 (UNC-16; UniProt entry P34609-1), kinesin heavy chain KIF5C (UNC-116; P34540-1), and dynein light intermediate chain (DLI-1; G5ED34-1). WT denotes wild-type. Molecular weight is indicated in kilodaltons (kDa). MBP::UNC-16[1-120] proteins (WT and V72Q) also contain a C-terminal Strep-tag II, which is detected on the immunoblot in *(A)* along with GST::DLI-1[369-442].

**Figure S4.**
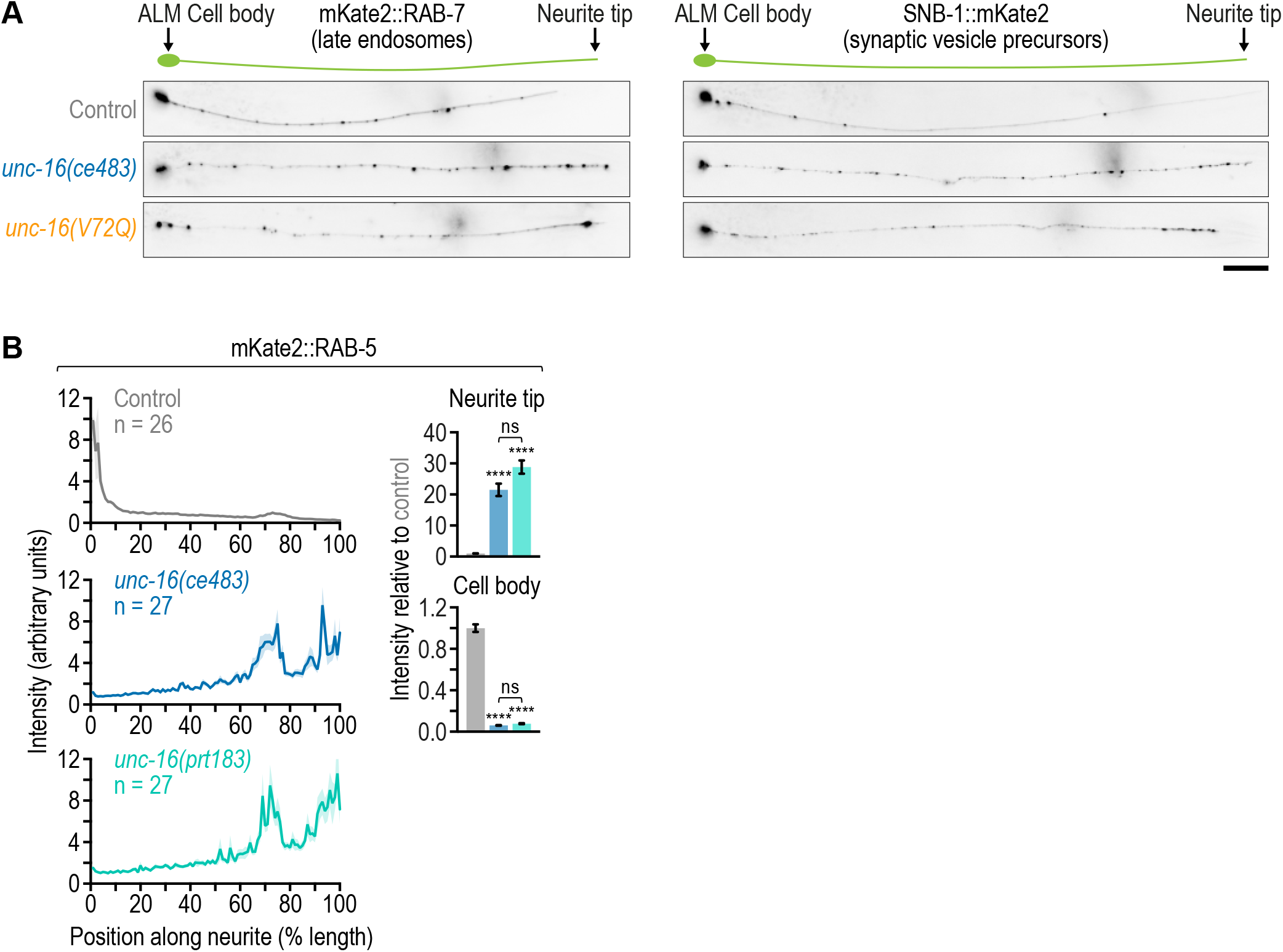
Late endosomes but not synaptic vesicle precursors are distributed differently in *unc/16(ce483)* versus *unc/16(V72Q)*d *unc/16(ce483)* is a null mutant. **(A)** Fluorescence images (maximum intensity z-stack projection, inverted grayscale) of the ALM neuron in L4 animals expressing a transgene-encoded marker for late endosomes (mKate2::RAB-7) or synaptic vesicle precursors (SNB-1::mKate2) in touch receptor neurons. Scale bar, 20 µm. **(B)** *(left)* Fluorescence intensity profiles (mean ± SEM) along the ALM neurite in L4 animals expressing a transgene-encoded marker for early endosomes (mKate2::RAB-5). *(right)* Integrated fluorescence intensity (mean ± SEM; normalized to control) in the ALM cell body and the last 20 µm of the distal neurite (neurite tip). *n* denotes the number of neurites examined. Statistical significance was determined by ANOVA on ranks (Kruskal-Wallis nonparametric test) followed by Dunn’s multiple comparison test. \*\*\*\**P* < 0.0001; *ns* = not significant, *P* > 0.05.

**Figure S5.**
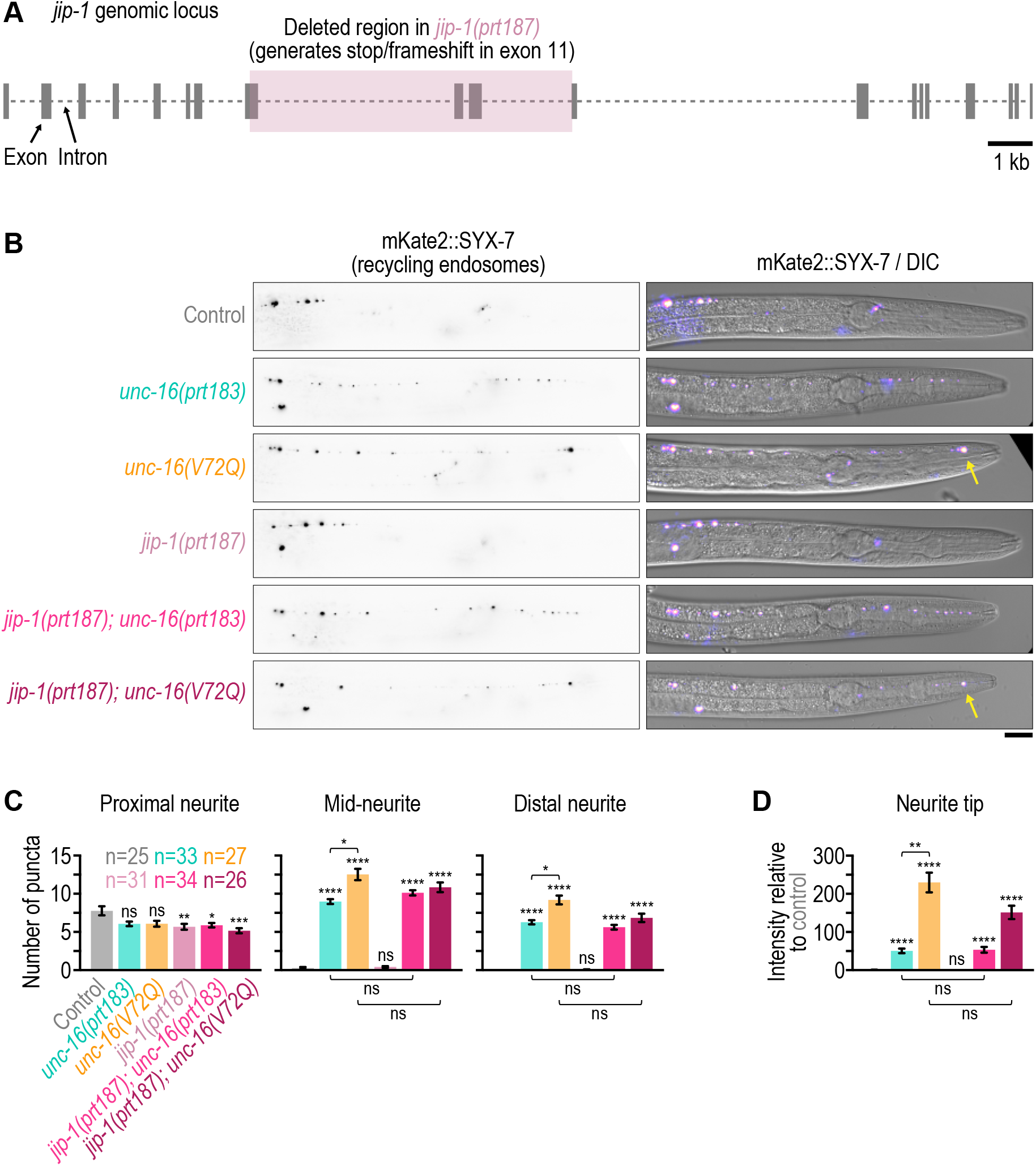
Recycling endosomes accumulate at the ALM neurite tip in the *unc/ 16(V72Q)* mutant and do so independently of JIP01. **(A)** Genomic locus of *jip:1* and modification in the *jip:1(prt187)* mutant introduced by genome editing. **(B)** Fluorescence images (maximum intensity z-stack projection, inverted grayscale) of the ALM neuron in L4 animals expressing a transgene-encoded marker for recycling endosomes (mKate2::SYX-7) in touch receptor neurons. Merged image on the right shows the location of the fluorescence signal relative to the differential interference contrast (DIC) image of the animal. Arrow points to the signal at the neurite tip. Note that diffuse signal in the neurite and the cell body is not readily discernable with this marker, which precludes determination of the fluorescence intensity profile along the neurite and intensity measurement in the cell body. Scale bar, 20 µm. **(C)** Number of mKate2::SYX-7 puncta (mean ± SEM) in the first quarter of ALM neurite length after the cell body (proximal neurite), the middle two quarters (mid-neurite), and the last quarter (distal neurite). *n* denotes the number of neurites examined (1 per animal). Statistical significance (control versus mutants; other comparisons indicated by brackets) was determined by ANOVA on ranks (Kruskal-Wallis nonparametric test) followed by Dunn’s multiple comparison test. \*\*\*\**P* < 0.0001; \*\*\**P* < 0.001; \*\**P* < 0.01; \**P* < 0.05; *ns* = not significant, *P* > 0.05. **(D)** Integrated fluorescence intensity (mean ± SEM; normalized to control) in the distal tip of the ALM neurite. The number of neurites examined (1 per animal) corresponds to the number *n* in *(C)*. Statistical significance was determined as described for *(C)*. \*\*\*\**P* < 0.0001; \*\**P* < 0.01; *ns* = not significant, *P* > 0.05.

**Table S1.** C. elegans strains.

| Strain | Genotype |
| --- | --- |
| N2 | Wild type (ancestral N2 Bristol) |
| GCP295 | *unc-119(ed3) III; prtSi101 [pRG526; mec-7p::snb-1::mKate2; cb-unc-119(+)] II* |
| GCP297 | *unc-119(ed3) III; prtSi104 [pRG527; mec-7p::mKate2::rab-5; cb-unc-119(+)] II* |
| GCP671 | *dli-1(prt106 [F392A/F393A]) IV/nT1 [qls51] (IV;V)* |
| GCP851 | *unc-119(ed3) III; prtSi166 [pRG528; mec-7p::mKate2::rab-7; cb-unc-119(+)] II* |
| GCP968 | *unc-16(ce483) III; prtSi104 [pRG527; mec-7p::mKate2::rab-5; cb-unc-119(+)] II* |
| GCP976 | *unc-16(prt162 [V72Q]) III* |
| GCP977 | *unc-16(prt162 [V72Q]) III; prtSi104 [pRG527; mec-7p::mKate2::rab-5; cb-unc-119(+)] II* |
| GCP978 | *unc-16(ce483) III; prtSi166 [pRG528; mec-7p::mKate2::rab-7; cb-unc-119(+)] II* |
| GCP979 | *unc-16(prt162 [V72Q]) III; unc-119(ed3) III; prtSi166 [pRG528; mec-7p::mKate2::rab-7; cb-unc-119(+)] II* |
| GCP1011 | *unc-16(prt162 [V72Q]) III; unc-116(e2310) III; prtSi104 [pRG527; mec-7p::mKate2::rab-5; cb-unc-119(+)] II* |
| GCP1012 | *unc-16(prt162 [V72Q]) III; unc-116(e2310) III; prtSi166 [pRG528; mec-7p::mKate2::rab-7; cb-unc-119(+)] II* |
| GCP1013 | *unc-119(ed3) III; prtSi212 [pRG1193; mec-4p::unc-16::egfp; cb-unc-119(+)] II* |
| GCP1025 | *unc-16(ce483) III; prtSi101 [pRG526; mec-7p::snb-1::mKate2; cb-unc-119(+)] II* |
| GCP1026 | *unc-16(prt162 [V72Q]) III; prtSi101 [pRG526; mec-7p::snb-1::mKate2; cb-unc-119(+)] II* |
| GCP1039 | *unc-119(ed3) III; prtSi215 [pRG1254; mec-7p::ctns-1::mKate2; cb-unc-119(+)] II* |
| GCP1044 | *unc-16(prt162 [V72Q]) III; prtSi209 [pRG1195; mec-4p::unc-16 (V72Q)::egfp; cb-unc-119(+)] II* |
| GCP1059 | *unc-16(prt162 [V72Q]) III; prtSi215 [pRG1254; mec-7p::ctns-1::mKate2; cb-unc-119(+)] II* |
| GCP1060 | *unc-16(ce483) III; prtSi215 [pRG1254; mec-7p::ctns-1::mKate2; cb-unc-119(+)] II* |
| GCP1061 | *unc-16(prt162 [V72Q]) III; unc-116(e2310) III; prtSi215 [pRG1254; mec-7p::ctns-1::mKate2; cb-unc-119(+)] II* |
| GCP1069 | *unc-16(prt177 [Δaa33-81]) III* |
| GCP1077 | *unc-16(prt177 [Δaa33-81]) III; prtSi104 [pRG527; mec-7p::mKate2::rab-5; cb-unc-119(+)] II* |
| GCP1088 | *dli-1(prt106 [F392A/F393A]) IV/nT1 [qls51] (IV;V); prtSi104 [pRG527; mec-7p::mKate2::rab-5; cb-unc-119(+)] II* |
| GCP1089 | *dli-1(prt106 [F392A/F393A]) IV/nT1 [qls51] (IV;V); prtSi212 [pRG1193; mec-4p::unc-16::egfp; cb-unc-119(+)] II* |
| GCP1111 | *unc-16(prt177 [Δaa33-81]) III; prtSi166 [pRG528; mec-7p::mKate2::rab-7; cb-unc-119(+)] II* |
| GCP1112 | *unc-16(prt177 [Δaa33-81]) III; prtSi215 [pRG1254; mec-7p::ctns-1::mKate2; cb-unc-119(+)] II* |
| GCP1115 | *unc-16(prt183 [ko]) III* |
| GCP1116 | *unc-16(prt183 [ko]) III; prtSi104 [pRG527; mec-7p::mKate2::rab-5; cb-unc-119(+)] II* |
| GCP1127 | *unc-119(ed3) III; prtSi236 [pRG1273; mec-7p::mKate2::syx-7; cb-unc-119(+)] I* |
| GCP1131 | *jnk-1(gk7) IV; prtSi104 [pRG527; mec-7p::mKate2::rab-5; cb-unc-119(+)] II* |
| GCP1132 | *jnk-1(gk7) IV; unc-16(prt162 [V72Q]) III; prtSi104 [pRG527; mec-7p::mKate2::rab-5; cb-unc-119(+)] II* |
| GCP1146 | *jip-1(prt187 [ko]) II; prtSi236 [pRG1273; mec-7p::mKate2::syx-7; cb-unc119(+)] I* |
| GCP1147 | *unc-16(prt162 [V72Q]) III; prtSi236 [pRG1273; mec-7p::mKate2::syx-7; cb-unc119(+)] I* |
| GCP1149 | *unc-16(prt183 [ko]) III; prtSi236 [pRG1273; mec-7p::mKate2::syx-7; cb-unc119(+)] I* |
| GCP1156 | *jip-1(prt187 [ko]) II; unc-16(prt162 [V72Q]) III; prtSi236 [pRG1273; mec-7p::mKate2::syx-7; cb-unc119(+)] I* |
| GCP1190 | *unc-16(prt190 [Δaa33-197]) III* |
| GCP1191 | *unc-16(prt190 [Δaa33-197]) III; prtSi104 [pRG527; mec-7p::mKate2::rab-5; cb-unc-119(+)] II* |
| GCP1192 | *unc-16(ce856 [L393P]) III; prtSi104 [pRG527; mec-7p::mKate2::rab-5; cb-unc-119(+)] II* |
| GCP1202 | *dli-1(prt106 [F392A/F393A]) IV/nT1 [qls51] (IV;V); prtSi215 [pRG1254; mec-7p::ctns-1::mKate2; cb-unc-119(+)] II* |
| GCP1206 | *unc-16(ce856 [L393P]) III; unc-16(prt191 [Δaa33-81]) III; prtSi104 [pRG527; mec-7p::mKate2::rab-5; cb-unc-119(+)] II* |
| GCP1234 | *unc-16(ce856 [L393P]) III; unc-16(prt205 [V72Q]) III; prtSi104 [pRG527; mec-7p::mKate2::rab-5; cb-unc-119(+)] II* |
| GCP1246 | *unc-16(prt183 [ko]) III; prtSi215 [pRG1254; mec-7p::ctns-1::mKate2; cb-unc-119(+)] II* |
| GCP1247 | *unc-16(ce856 [L393P]) III; prtSi215 [pRG1254; mec-7p::ctns-1::mKate2; cb-unc-119(+)] II* |
| GCP1248 | *unc-16(ce856 [L393P]) III; unc-16(prt205 [V72Q]) III; prtSi215 [pRG1254; mec-7p::ctns-1::mKate2; cb-unc-119(+)] II* |
| FF41 | *unc-116(e2310) III* |
| KG2338 | *unc-16(ce483) III* |
| KG5309 | *unc-16(ce856 [L393P]) III* |
| VC8 | *jnk-1(gk7) IV* |

**Table S2.**
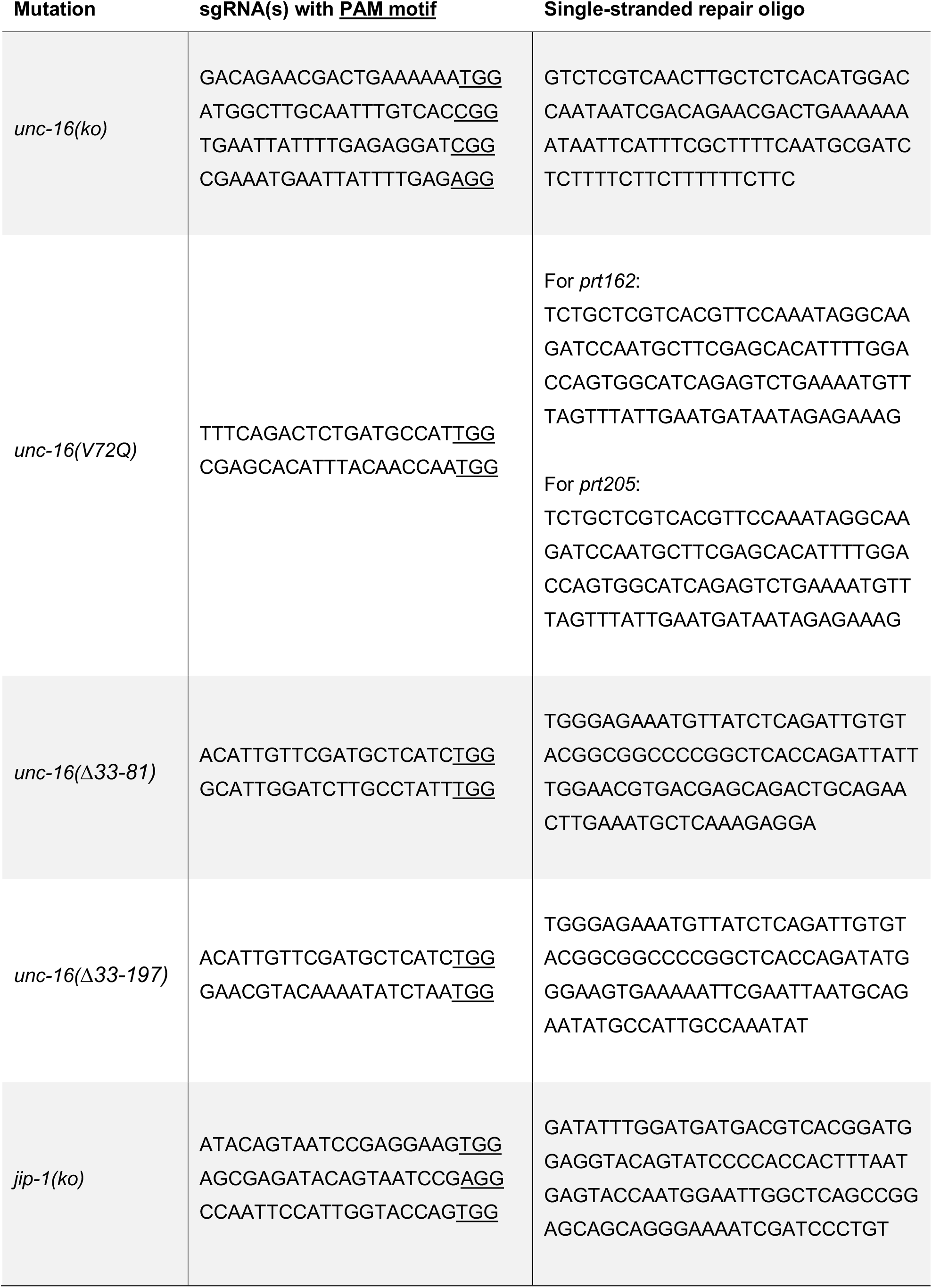
Oligonucleotides for CRISPR/Cas9@mediated genome editing.

